# Modelling the spatiotemporal dynamics of senescent cells in wound healing, chronic wounds, and fibrosis

**DOI:** 10.1101/2024.07.04.602041

**Authors:** Sharmilla Chandrasegaran, James P. Sluka, Daryl Shanley

## Abstract

Cellular senescence is known to drive age-related pathology through the senescence-associated secretory phenotype (SASP). However, it also plays important physiological roles such as cancer suppression, embryogenesis and wound healing. Wound healing is a tightly regulated process which when disrupted results in conditions such as fibrosis and chronic wounds. Senescent cells appear during the proliferation phase of the healing process where the SASP is involved in maintaining tissue homeostasis after damage. Interestingly, SASP composition and functionality was recently found to be temporally regulated, with distinct SASP profiles involved: a fibrogenic, followed by a fibrolytic SASP, which could have important implications for the role of senescent cells in wound healing. Given the number of factors at play a full understanding requires addressing the multiple levels of complexity, pertaining to the various cell behaviours, individually followed by investigating the interactions and influence each of these elements have on each other and the system as a whole. Here, a systems biology approach was adopted whereby a multi-scale model of wound healing that includes the dynamics of senescent cell behaviour and corresponding SASP composition within the wound microenvironment was developed. The model was built using the software CompuCell3D, which is based on a Cellular Potts modelling framework. We used an existing body of data on healthy wound healing to calibrate the model and validation was done on known disease conditions. The model provides understanding of the spatiotemporal dynamics of different senescent cell phenotypes and the roles they play within the wound healing process. The model also shows how an overall disruption of tissue-level coordination due to age-related changes results in different disease states including fibrosis and chronic wounds. Further specific data to increase model confidence could be used to explore senolytic treatments in wound disorders.

## Introduction

Cellular senescence is a process whereby cells undergo an irreversible cell cycle arrest in response to a diverse range of stresses. It was first discovered by Hayflick and Moorhead, who demonstrated that cultured human fibroblasts had an exhaustive replicative capacity before entering a state of replicative senescence (RS) [1]. The irreversible cell cycle arrest in senescence is also accompanied by distinct phenotypic and metabolic features, as well as the senescence-associated secretory phenotype (SASP). SASP is characterised by the transcriptional activation of several factors including growth factors, cytokines, chemokines, extracellular matrix (ECM) components and matrix proteinases [2].

Senescent cells can exert effects on their surrounding microenvironment in many ways depending on context. They accumulate in tissues and organs with age and, as such, senescence is one of the hallmarks of ageing [3]. Senescent cells are involved in many age-related pathologies including Parkinson’s disease [4], osteoarthritis [5], atherosclerosis [6], fibrosis [7], chronic wounds [8] and many others [9]. However, they also play many beneficial roles, primarily in the context of embryonic development [10, 11], tumour suppression [12–14], and wound healing and tissue repair [7, 8, 15].

During wound healing, senescent cells transiently appear during the mid to late stages of tissue repair and are removed by the immune system during the later stages. Several SASP factors, including PDGF and MMPs, play important roles in maintaining tissue homeostasis during the repair process [16]. For instance, senescent cells have been shown to mitigate fibrosis during tissue repair in the skin, cornea, liver, heart and pancreas [17–21]. Aside from being an anti-fibrotic mechanism, they also promote myofibroblast differentiation through SASP, which is crucial for ECM production and dermal wound closure [15]. However, senescence has also been implicated in the pathophysiology of fibrotic conditions across several organ systems [22–24], and the accumulation of senescent cells has been linked to the development of chronic wounds [25–27]. This is interesting since fibrosis and chronic wounds represent two extreme situations equivalent to ‘over-healed’ and ‘under-healed’ wounds respectively.

Beneficial effects of senescent cells are evident from their transient role in tissue repair, with the absence of senescent cells impairing the repair process, promoting pro-fibrotic activity and tumorigenesis [16–21]. In contrast, an excess of senescent cells results in unresolved inflammation [25, 28, 29]. Persistent senescent cells, despite cell cycle arrest, remain metabolically active within the wound microenvironment and can communicate with surrounding cells through paracrine signalling. This is further complicated by the composition and functionality of SASP produced within the repair system which has been shown to be temporally regulated by Notch signalling; the beginning of the senescence programme involves a fibrogenic SASP, followed by a fibrolytic SASP [30–32]. Keeping in mind that tissue repair is an inherently complex process, with several other factors at play, the timing of induction of senescent cells and variations in SASP composition during wound repair could steer the process towards different outcomes, either healthy repair or towards fibrosis or chronic inflammation. The interplay between varying senescent cell activity and their surroundings could lead to a wide range of qualitatively distinct spatiotemporal dynamics, that could result in these different repair outcomes.

To investigate this, we present a multi-scale computational model of wound healing that includes the dynamics of senescent cells. This model is used to demonstrate and investigate the range of senescent cell functional heterogeneity within dysregulated repair, namely, fibrotic, and chronic wounds.

## Model and Methods

The wound healing model was built using the software CompuCell3D (CompuCell3D.org)[33], a modelling environment implementing the lattice-based Cellular Potts Model (CPM) framework, with the aim of investigating the spatiotemporal dynamics of senescent cells in wound healing, fibrosis, and chronic wounds [33]. Given the complex spatial and multiscale nature of the processes involved in wound healing and tissue repair, the CPM framework was used since it explicitly represents space and can combine processes occurring at various length and time scales. To develop our mechanism-based computational model, biological observations were gathered to formulate a conceptual model, as summarised in Figure 1. Once completed, a qualitative model was built including the essential model elements that capture the observed biology, along with abstractions and assumptions using the CPM framework.

**Figure 1.**
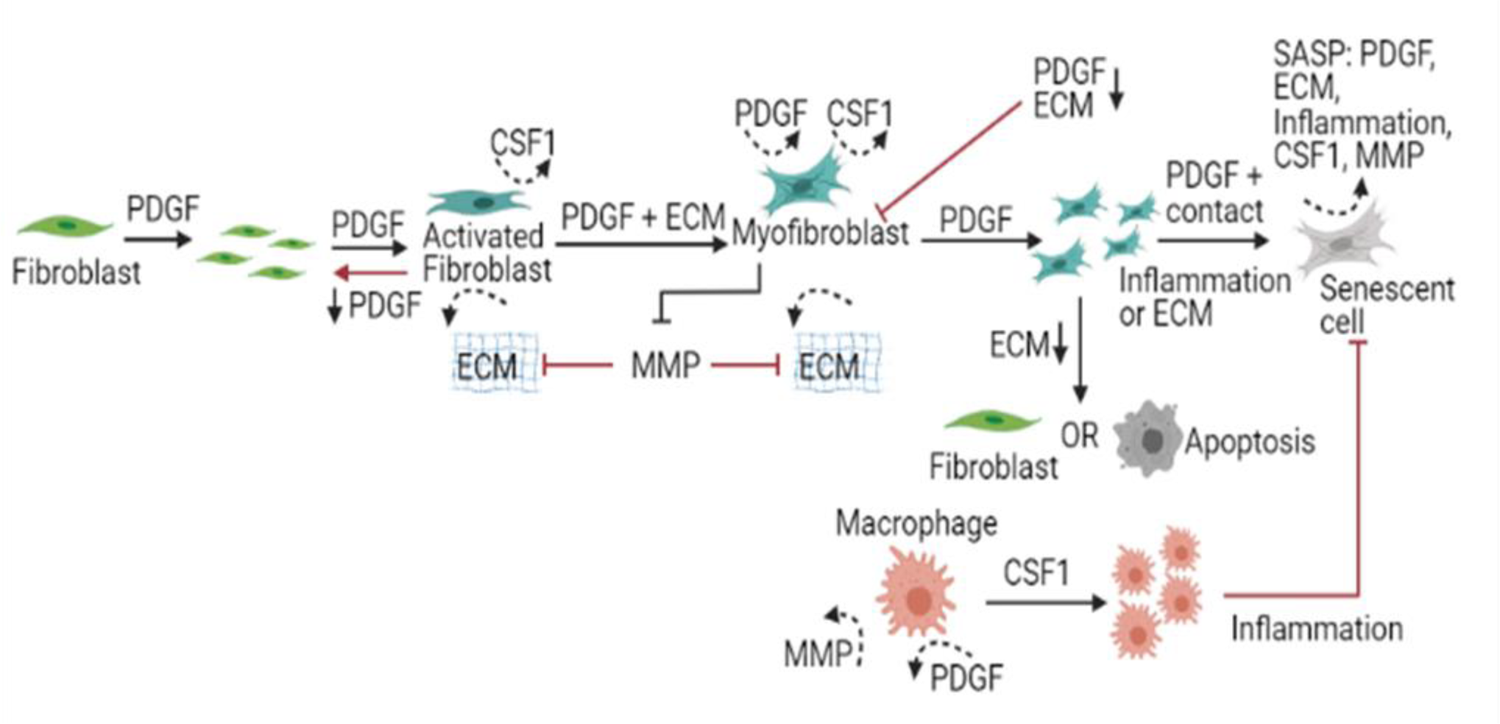
Conceptual illustration of the wound healing model. This is a schematic representation of the qualitative model which was developed before being applied to a computational framework. The model consists of explicitly described generalised cell types including fibroblasts, myofibroblasts, senescent myofibroblasts, macrophages and ECM. Chemical fields within the model include PDGF, CSF1, MMP and Inflammation. Fibroblasts proliferate and become activated in response to PDGF. Activated fibroblasts produce ECM and CSF1. In response to PDGF and mechanical stimuli, activated fibroblasts differentiate into myofibroblasts, which produce large amounts of ECM, PDGF and CSF1. Myofibroblasts are highly proliferative cells that replicate in response to PDGF. Myofibroblasts become senescent through CCN1-induced primary senescence, and paracrine- and juxtacrine-induced secondary senescence. Macrophages on the other hand proliferate in response to CSF1 and are the primary source of several growth factors within the wound environment.

In the CPM framework, the simulation lattice is divided into collections of pixels representing generalised ‘cells’, each with an assigned ‘cell’ type. ‘Cells’ interact with each other and their environment based on parameters that can approximate biological constraints. These parameters can be based on their *in vitro* or *in vivo* biological counterparts and steer the simulation. The CPM evolves through time using an effective energy Hamiltonian (H) and Boltzmann Factor acceptance function that is calculated at each simulation time step (a Monte-Carlo step, MCS). The Boltzmann Factor uses a Potts “Cell membrane fluctuation amplitude” (T_m_) that allows the simulated cells to explore an energetically plausible trajectory through time instead of simply minimizing the calculated system energy, which tends to get trapped in local energy minima. Terms of the effective energy equation for the wound healing model will be described in the next sections. Model parameter estimation and detailed simulation specifications are described in the Supplementary methods. The source CC3D code is also available.

The multiscale model of wound healing represents a two-dimensional wounded dermal section that is of size 200×200×1 pixel (600 µm × 600 µm). Pixel units and corresponding cell lengths and volumes are provided in Supplementary Table 1. All cells are chemically homogeneous as no chemical diffusion is modelled within the intracellular space; this was to keep the focus on tissue-level and intercell behaviours. Since the primary focus of the model is to investigate the dynamics of senescent cells in the healing process, the model simulation begins at the end of the inflammatory phase, since the proliferative phase is when senescent cells are most prevalent in healthy wound healing [16]. Furthermore, the papillary and reticular dermis are assumed to be combined within the model, similar to previous work from Rognoni et al. where papillary and reticular fibroblast populations were represented by a single cell type for the purpose of modelling simplification [34].

### Cell dynamics and motility

Cell motility driven by the cytoskeleton is modelled within the CPM using a modified Metropolis algorithm which consists of a series of stochastic pixel-copy attempts, described in more detail in Supplementary methods (Cell dynamics and motility) [33, 35].

One pixel in the simulation was taken to be equal to 3µm, as this seemed to be a reasonable trade of between cellular spatial detail and computational speed. Given this pixel dimension, a typical cell in the simulation is 8×8 pixels for an area or 200um^2^. Cells in the simulation were found to have an average speed of ∼0.1 pixel/MCS, which equated to 0.2 microns/MCS. This speed was determined by running multiple simulations of a cell without any hindrance and taking an average of the difference between the current and previous center of mass (COM) values. To calculate the real time equivalent of one MCS, maximum fibroblast migration speed during wound healing was considered as a reference, which peaks towards the end of the healing process with a speed of 40 µm/h [36]. Therefore, one MCS was calculated to be 27 secs (0.3*3600/40).

### Cell types, volume, surface and adhesion

The model consists of fibroblasts, myofibroblasts, macrophages, senescent myofibroblasts and ECM defined as individual cell types with distinct properties, as shown in Figure 1. The dermal layer is composed of ECM and resident fibroblasts, whereas the wound is populated by M2-polarised macrophages. Fibroblasts differentiate into myofibroblasts through mechanisms described in later sections. Myofibroblasts can either undergo apoptosis or become senescent myofibroblasts. Terms in the effective energy equation describing cell volume and surface, and cell adhesion are presented in the Supplementary methods.

### Cell growth and proliferation

Within the model, cell growth is limited to fibroblasts, myofibroblasts and macrophages. As discussed previously, PDGF is a primary mediator of fibroblast and myofibroblast migration, proliferation, and ECM production within the wound [37, 38]. Hence, fibroblast and myofibroblast growth were modelled by increasing cell target volume according to PDGF concentration using the Monod growth equation given as,

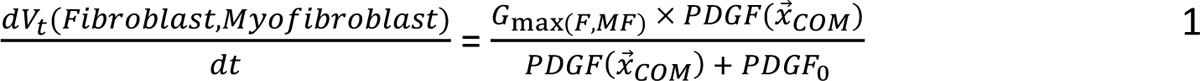

Where *PDGF*(*x*→_*COM*_) is PDGF concentration at the cell’s COM, *PDGF*_0_ is the concentration at which cell growth is half its maximum, and *G*_max(*F,MF*)_ is the maximum cell cycle time for fibroblasts and myofibroblasts. Similarly, CSF1-dependent macrophage growth is given as,

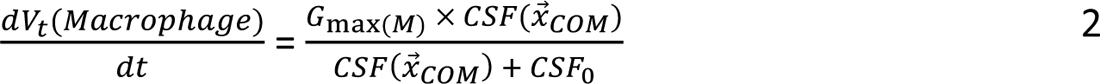

Where *CSF*(*x*→_*COM*_) is CSF1 concentration at the cell’s COM, *CSF*_0_ is the concentration at which growth is half its maximum, and *G*_*max*(*M*)_ is the maximum cell cycle time for macrophages. The Monod equation relates cell growth to the concentration of a limiting factor, which in this case is PDGF for fibroblasts and myofibroblasts, and CSF1 for macrophages [39].

Cell growth and proliferation are limited by contact inhibition, which can occur through two ways: mechanical stress and contact-dependent growth inhibition [40, 41]. Mechanical stress-dependent inhibition was modelled by imposing a volume constraint for mitosis whereby cells undergo mitosis only when they reach their doubling volume and then divide along a randomly chosen axis. Cell doubling volume was taken as twice the original target volume of that cell type (fibroblasts, myofibroblasts or macrophages), with the daughter cells acquiring the target volume of the parent cells.

Contact-dependent growth inhibition was modelled by allowing an increase in cell volume only if the surface of the cell in contact with neighbouring cells, *R*_*s*_, is below a specified threshold *T*_*CI*_ [42]. *R*_*s*_ is given by:

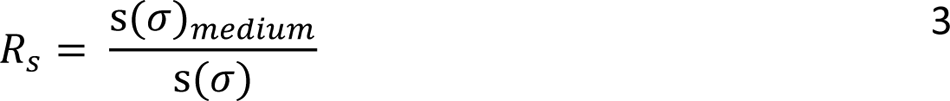

Where s(σ)_*medium*_ is the cell surface area in contact with the medium and s(σ) is the cell surface area. The ratio *R*_*s*_ measures how much of a cell’s surface area is not in contact with other neighbouring cells. If *R*_*s*_ is below the threshold, *R*_*CI*_ then cell growth is arrested at the current cell volume *v*(σ) along with the initial lambda value λ_*vol*_(σ), and if *R*_*s*_ is above *R*_*CI*_, cell volume is determined by equations 1 and 2.

### Cell state transitions, growth factors and cytokines

Fibroblasts, myofibroblasts and macrophages interact by exchanging growth factors. Macrophages respond to various external stimuli that can result in a wide range of activation phenotypes [43]. Macrophages within the model shift between classically (M1) and alternatively (M2) activated phenotypes depending on contact with the inflammation field, which represents inflammatory SASP within the model [44]. The inflammatory component of SASP is highly complex and consists of many key factors [32]. However, for the sake of computational feasibility, inflammatory SASP was considered as one chemical field.

If the concentration of the inflammation field at the COM of a macrophage is greater than the threshold parameter *INF_thr_*, then the macrophage secretes MMPs (also referred to as proteinase), characteristic of the M1 phenotype. However, if the inflammation field concentration is less than *INF_thr_*, the macrophage secretes PDGF [45]. MMPs can also be produced by macrophages following contact with ECM, that is, if ECM is surrounding the cell periphery [46]. Since the model begins at the end of the inflammation phase, macrophages are M2 activated and secrete PDGF for fibroblasts and myofibroblasts [47]. For the sake of simplicity, and to focus on senescent cell activity during the proliferative phase of healing, inflammatory SASP is the only form of inflammation considered in this model.

Quiescent fibroblasts become activated in response to PDGF, stimulating the production of CSF1 [48, 49]. Myofibroblasts also produce CSF1 for macrophages, as well as PDGF, forming an autocrine loop [50]. In the model, if PDGF concentration at a fibroblast’s COM is above the threshold parameter *PDGF_f_*, then the fibroblast is activated from a quiescent state. Growth factors, especially CSF1, exchange occurs in a cell-cell contact-dependent manner [51]. Therefore, CSF1 secretion by activated fibroblasts and myofibroblasts only occur in cells sharing common surface area with macrophages. Activated fibroblasts and macrophages in the model were removed at a rate of µ[51]. Macrophages also undergo apoptosis following growth factor withdrawal [52, 53], which was modelled by removing macrophages with CSF1 concentration less than a threshold parameter *CSF1_thr_*at the cell’s COM.

Myofibroblast differentiation typically requires growth factor stimulation, along with mechanical stimulation [54]. PDGF is one growth factor that is known to play an important role in granulation tissue formation and myofibroblast differentiation during tissue repair [55–57]. In addition, mechanical stimulation is crucial for the formation of the contractile features characteristic of myofibroblasts [58–60]. In the model, myofibroblast differentiation occurs if the value of PDGF at a fibroblast’s COM is greater than *PDGF_f_*and common surface area with ECM is greater than *ECM_thr_*. Every activated fibroblast within the wound region in the model has a chance of differentiation that is calculated by drawing a random integer from a binomial distribution with probability *P_MF._* Since myofibroblasts are known to produce TIMPs, a simple mechanism to inhibit proteases was included, whereby MMP concentration at a myofibroblast’s COM was reduced by the MMP threshold *MMP_thr_* [61]. An important feature of fibroblasts and myofibroblasts is their ability to produce large amounts of ECM. Fibroblasts and myofibroblasts produce ECM if PDGF concentration at the cell’s COM is above the threshold parameter *PDGF_f_*. The amount of ECM produced by myofibroblasts (*ECM_Myof_*) is assumed to be double that produced by fibroblasts (*ECM_Fib_*). This value was selected to be most appropriate after running multiple preliminary model simulations with different parameter values. ECM degradation by MMP occurs by removing ECM ‘cells’ that have a MMP concentration at their COM greater than *MMP_thr_*.

Myofibroblast activity can resolve in many ways. Myofibroblasts can undergo apoptosis due to release of mechanical tension or exposure to pro-apoptotic inflammatory cytokines, de-differentiate to a fibroblast phenotype or become senescent [62]. To model apoptosis due to release of mechanical tension, myofibroblasts that shared common surface area with ECM below a threshold represented by the parameter *ECM_thr_*, were removed [63]. Apoptosis by exposure to inflammation was modelled by removing a cell if concentration of the inflammatory SASP field at the COM of the cell was greater than the threshold parameter *INF_thr_* [64]. The chance of a myofibroblast becoming senescent was calculated by drawing a random integer from a binomial distribution with probability *P_SNC_*. Senescence can be induced in myofibroblasts through the matricellular protein CCN1 during the mid to late stages of healing [15, 16, 20]. Therefore, a time constraint *T_SEN_* was set for the induction of senescence to limit it to the mid to late stages of the repair process. In keeping with this, if time is less than *T_SEN_*, a myofibroblast in the model with common surface area with ECM that is greater than the parameter *ECM_thr_* will become a senescent myofibroblast. Senescent myofibroblasts produce two distinct types of SASP: a fibrogenic SASP consisting of PDGF and ECM lasting for about three days (*T_NIS_*), followed by a fibrolytic SASP consisting of CSF1, Inflammation and MMP. This value for *T_NIS_* was chosen because preliminary work suggests that NOTCH-mediated fibrogenic SASP and juxtacrine secondary senescence is prevalent up to approximately three days after senescence activation [32, 65].

In addition to CCN1-induced senescence, mechanisms of secondary senescence include juxtacrine and paracrine senescence, which are more prevalent during the fibrogenic and fibrolytic phases of the senescence program respectively [32]. Therefore, within the model, the juxtacrine mechanism involves induction of senescence if a myofibroblast is in contact with a senescent cell in its fibrogenic SASP phase, and if PDGF concentration at the myofibroblast’s COM was greater than the threshold parameter *SNC_thr_*. For paracrine senescence, if a myofibroblast is neighbouring a senescent cell in its fibrolytic SASP phase and had an inflammatory SASP field concentration greater than *INF_thr,_* then the cell would become senescent. Senescent cells during wound healing are removed by immune cells, e.g., phagocytosis by macrophages [66]. Hence, senescent cells within the model are removed on contact with macrophages to model phagocytosis. They also have an equal chance of getting removed if the concentration of the inflammation field at the senescent cell’s COM is greater than *INF_thr_*, to account for other immune-surveillance mechanisms.

The governing reaction-diffusion equations for the four chemical fields in the model are given as below:

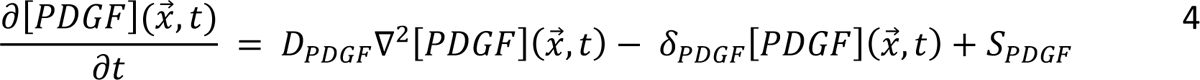

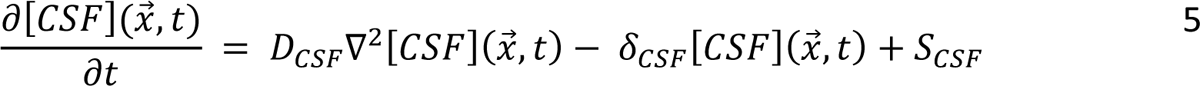

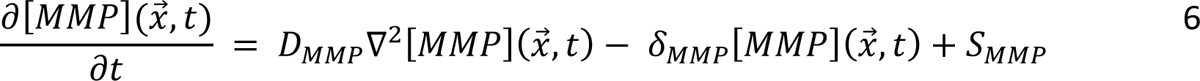

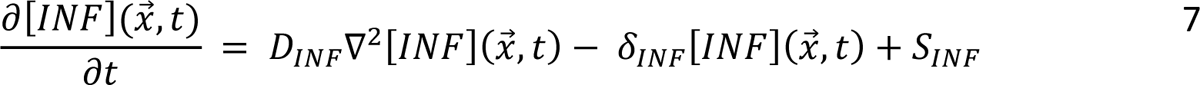

Where [*PDGF*](*x*→, *t*), [*CSF*](*x*→, *t*), [*MMP*](*x*→, *t*), [*INF*](*x*→, *t*) represent the concentrations of the chemical fields at pixel *x*→ at time *t*, *D* represents the diffusion coefficients, δ represents the decay constant and *S* represents the rate of secretion for the fields PDGF, CSF1, INF and MMP. All chemical fields were set to no flux in the x and y axes. The solver ‘DiffusionSolverFE’ within CompuCell3D was used to solve the PDEs. Note that for simplicity, and to correct for the limitations of a 2D model, diffusible species are modelled as if they can diffuse across cells.

### Chemotaxis

Chemotaxis refers to cell migration directed towards a chemical stimulus and is an important element in wound healing as it is one of the main mechanisms by which cells are recruited to the wound site. Within the model, chemical field gradients of PDGF and inflammation cause cells to chemotax towards them. Both fibroblast and myofibroblast chemotaxis towards PDGF [67]. Fibroblast migration is directed by orientation of ECM fibres and they travel along the ECM through integrin-collagen adhesions [68]. To model this effect of ECM fibres on cell migration, the chemotaxis strength coefficient was increased to λ_*PDGF*(*ECM*)_ for fibroblasts and myofibroblasts in contact with ECM. Macrophages, which are involved in immune surveillance for senescent cells, chemotax towards the inflammation field produced by senescent cells [69]. Terms in the effective energy equation describing cell chemotaxis are detailed in Supplementary methods.

### Model classifications

To investigate the spatiotemporal dynamics of senescent cells during tissue repair, perturbations to senescent cell dynamics within the model were introduced. Additionally, another model initial state including pre-existing inflammatory senescent cells was created to represent both age-related changes, and the early induction of senescent cells during the inflammatory phase of healing. These inflammatory senescent cells produce inflammatory SASP, CSF1 and proteinase within the model, with all other properties (such as size and contact energies) the same as for senescent myofibroblasts.

To study the changes induced by differences in senescent cell activity alone, no other factors or parameters were changed in this investigation. Two parameters related to senescent cell dynamics in the model were changed to produce the different outcome classifications: The probability of senescence in myofibroblasts, which is dictated by the parameter *P_SNC_* and the time of induction of senescent cells in the model, determined by the time constraint parameter *T_SEN_*. Simply put, the parameter *P_SNC_* controls the population size of senescent cells, and the parameter *T_SEN_* controls when senescent cells are induced at the wound site. Original model values are 0.15 for *P_SNC_* and 12 for *T_SEN_* (see Supplementary Table 1).

To investigate changes in wound healing in response to a lack of senescent cells, *P_SNC_* was set to zero, with no changes to *T_SEN_*. Conversely, to investigate the effects of increased number of senescent cells within the wound, *P_SNC,_* was set to 0.75, with no changes to *T_SEN_*.

To explore the temporal aspects of senescence, two scenarios were considered. The first scenario was the early induction of senescence during the beginning of the inflammatory phase (representing age-related accumulation of senescent cells), for which the model with pre-existing inflammatory senescent cells was used with no other changes. The second scenario was the late induction of senescent cells during the later phases of healing; in this case, for the model, *P_SNC_* was not altered, but *T_SEN_*was set to 15 days. Three replicate runs were performed for each classification.

### Multidimensional sensitivity analysis

To investigate the trade-offs between the two parameters, probability of senescence in myofibroblasts *P_SNC_* and time of senescence induction *T_SEN_*, a multidimensional sensitivity analysis was performed using the parameter sweep feature within CompuCell3D. Parameter value ranges were chosen as evenly spaced values within a defined range. Simulation time series data was collected every 500 MCS (model time step; equal to 3.75 hours) for each parameter set considered. A total of 25 simulations were run.

### Particle swarm optimisation parameter search

The wound healing model has many parameters, some of which are not available in the literature. To determine these parameters, we used a Particle swarm Optimization (PSO) approach[70]. Briefly, a PSO treats a particular set of parameters as a point (particle) in a multidimensional parameter space. For each particle, a quality metric is calculated using the model. The PSO generate multiple such parameter particles that propagate through parameter space looking for an optimal solution. Individual particles change their values based on the best result they have encountered as well as the best point encountered by the entire swarm. A swarm of particles will coalesce on a particular set of parameters in the parameter space, which may or may not be a global minimum. Multiple independent swarms are used to ensure adequate sampling of the entire parameter space. If multiple swarms find the same set of parameters, then that suggests the solution is either a global minimum, or at least a local minimum that is likely to be similar in quality to the (unknown) global minima.

The metric we used to describe the quality of a particular set or parameters was determined using relative error between simulation output and data from literature where possible. This includes cell population profiles for fibroblasts, myofibroblasts, macrophages and senescent myofibroblasts, ECM production and rate of wound closure, which are shown in Supplementary Figure 1 A-F.

The parameters within the wound healing model that we varied are fibroblast activation PDGF threshold (*PDGF_f_)*, PDGF concentration threshold for myofibroblast juxtacrine senescence (*SNC_thr_*), proteinase threshold for ECM breakdown (*MMP_thr_*), myofibroblast deactivation ECM threshold (*ECM_thr_*), Inflammation threshold (*INF_thr_*) and CSF1 threshold for macrophage depletion (*CSF1_thr_*) (Supplementary Table 2). For more information see the supplemental material.

## Results

### A multiscale model of healthy wound healing including senescent cell dynamics

The multiscale model of healthy wound healing developed here includes cells, growth factors and cytokines crucial for the wound healing process. In contrast to existing computational models of wound healing, this model includes the dynamics of senescent cells and key SASP components involved in the healing process.

Briefly, cell types within the model were represented by collections of pixels with unique identifiers. Cells interact with each other based on parameters that were obtained from literature or estimated. The parameters steer model behaviour through the effective energy or Hamiltonian equation, with the simulation evolving stochastically through a Monte Carlo with a Boltzmann acceptance function, described in more detail in the Supplementary results. Growth factors and cytokines within the model were defined using reaction diffusion PDEs.

The wound healing model had a large number of parameters, some of which were obtained from literature, summarised in Supplementary Table 1, whereas others were estimated using PSO. The best parameter sets identified by PSO are given in Supplementary Table 2. Summary of the output from running PSO is in the Supplementary Table 3. PSO was run for a total of 60 iterations, and the best parameter set was found at the 46^th^ iteration.

Parameter values from swarm 1 were used for the final model as it had a slightly better fitness measure (relative error of 27.84). Plots of the resulting simulation data, plotted against data used for calibration for fibroblasts, senescent fibroblasts, myofibroblasts, ECM, macrophages and wound closure rate are shown in Figure 3A-F, along with the simulation cell field in Figure 2 showing a successfully healed wound. Despite the effort put into making the model parameters as physiologically and biologically relevant as possible, it was not possible to match or find appropriate data for some parameters. For these, parameter values were simulated from theoretically sound estimates to produce a biologically reasonable representation of the wound healing process based on information gathered from literature, as previously stated, and provided in Supplementary Table 1.

**Figure 2.**
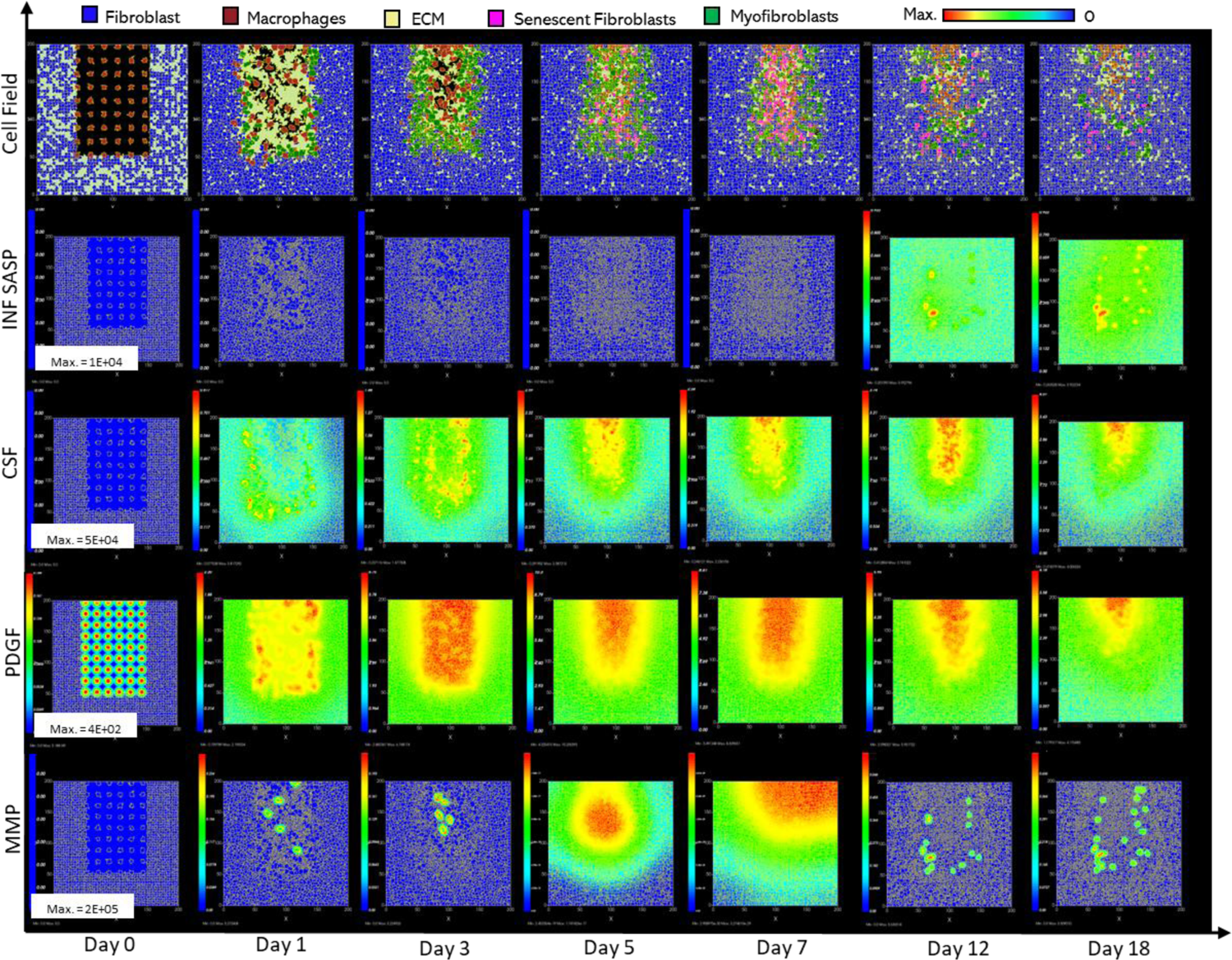
Simulation of healthy physiological wound healing in a dermal patch of size 600 µm x 600 µm using the baseline parameters. This figure shows snapshots of spatial configuration *vs* time for the progression of a simulated wound healing process. The simulation represents healthy wound healing from the end of the inflammatory phase to the remodelling phase, including fibroblasts, myofibroblasts, macrophages, ECM and senescent myofibroblasts, and chemical species including inflammatory SASP, CSF1, PDGF and MMP. **Top row:** Snapshots of the simulation cell field at different time points with fibroblasts (blue), myofibroblasts (green), macrophages (brown), ECM (yellow) and senescent myofibroblasts (pink). Snapshots of simulation chemical fields are shown from the second to the last row; **Second row:** Inflammatory SASP, **Third row:** CSF1, **Fourth row:** PDGF, **Fifth row:** MMP. Fields are shaded where, red corresponds to the maximum value specified in the first snapshot panel for each field, and blue corresponds to zero (shown in the colour bar at the top). Snapshots are shown for the time points, left to right, day 0, 1, 3, 5, 7, 12 and 18.

**Figure 3.**
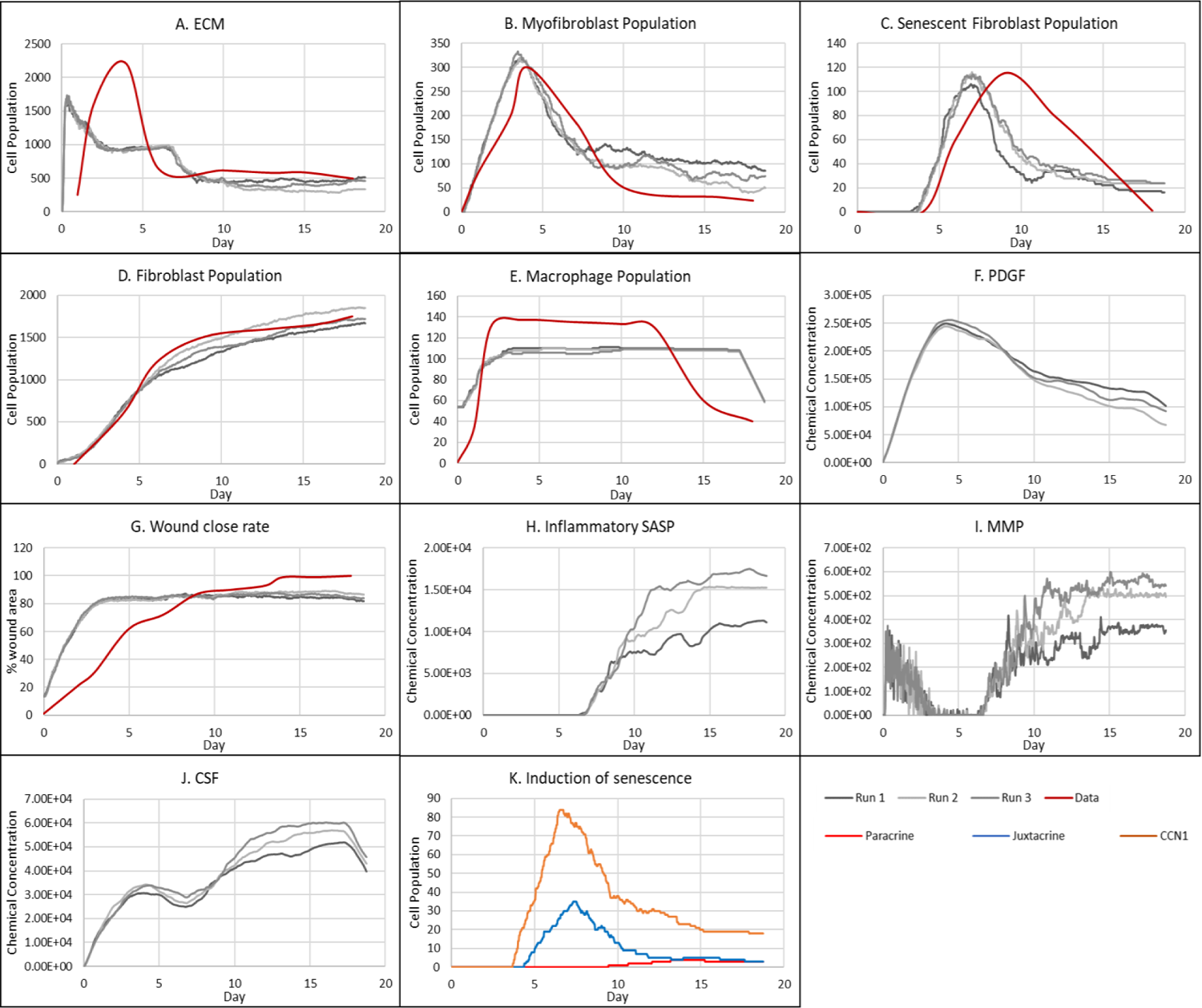
Simulation time series for the model of healthy physiological wound healing in a dermal patch of size 600 µm x 600 µm using the baseline parameters presented in Supplementary Table 1, shown for model cell types, rate of wound closure and model chemical fields. This figure shows the simulation time series for the progression of a simulated wound healing process, with number of cells or rate of wound closure on the y axis and time (day) on the x axis, shown for **A:** ECM, **B:** Myofibroblasts, **C:** Senescent myofibroblasts, **D:** Fibroblast, **E:** Macrophages, **F:** PDGF, **G:** Wound closure rate, **H:** Inflammatory SASP, **I:** MMP, **J:** CSF1 and **K:** Induction of senescence. For the time series plots A-F, the three shades of grey represent the different replicate runs as indicated below plot E (since the model is stochastic); the red line represents data used for parameter estimation using PSO informed by existing studies, as shown in Supplementary Figure 1. For the plot G, different shades of grey represent the different mechanisms of senescence induction, as indicated below the plot; dark grey: paracrine secondary senescence; light grey: juxtacrine secondary senescence; and medium grey: CCN1-induced primary senescence. Replicate runs are not shown for the different mechanisms for ease of readability, but total senescent cell numbers for the three replicate runs are shown in plot B.

Healthy wound healing is accompanied by tightly controlled transient senescent cell activity The model begins with a dermal layer consisting of fibroblasts and ECM along with a wound populated by anti-inflammatory M2 macrophages, representative of the end stages of the inflammatory phase of wound healing (Figure 2). Time series profiles for selected factors in the proliferative phase are shown in Figure 3. Anti-inflammatory M2 polarised macrophages are prevalent during the proliferative phase and are an important source of PDGF for fibroblasts. As a result, PDGF levels begin to rise steadily until approximately ∼day 4 (Figure 3F), setting off a cascade of events that will eventually result in myofibroblast differentiation, which will be described hereafter. As the simulation progresses, fibroblasts proliferate in response to rising PDGF levels resulting in a steady increase in fibroblast numbers at around ∼day 1 to day 5 after which it rises at a slower pace, corresponding to PDGF levels, which begin to subside at near ∼day 4 (Figure 3D and F).

PDGF also induces fibroblast activation, which leads to the production of CSF1 and ECM. This leads to rising CSF1 levels (Figure 3J), which results in macrophage proliferation and an increase in macrophage numbers by 2-fold, before reaching a plateau at ∼day 3.5 (Figure 3E). Macrophage numbers correspond to CSF1 levels, which begin to drop briefly around the same time (Figure 3J). Levels of ECM rise sharply at day 1, which is a simple representation of Type III collagen and FN-rich granulation tissue (Figure 3A). The production of ECM at this point, during the transition from inflammatory to proliferative phase, is an important source of mechanical stimulation which is necessary for myofibroblast differentiation of fibroblasts.

Appropriate PDGF levels and mechanical stimuli promote full myofibroblast differentiation of the activated fibroblasts. Myofibroblast numbers begin to rise at day 1, peaking at day 4 (Figure 3B). Myofibroblasts also produce large amounts of ECM, along with PDGF, in an autocrine manner, as well as CSF1 for macrophages. However, ECM levels plateau after a 0.7-fold drop at ∼day 3 during this rise in myofibroblast levels, due to macrophage-derived MMP, which breaks down excess ECM (Figure 3E and I). MMP levels also experience a decline due to its enzymatic activity on ECM and inhibition by myofibroblasts through TIMP activity. The plateau in ECM levels demonstrates a balance between ECM production by myofibroblasts and fibroblasts, and breakdown by MMPs. Myofibroblast numbers eventually begin to decline around day 4 by 0.5-fold, mediated by either apoptosis (due to release of mechanical tension), de-differentiation to a fibroblast phenotype or cellular senescence (Figure 3B).

Around the same time as the drop in myofibroblast numbers, the number of senescent myofibroblasts begins to rise at ∼day 4, reaching a peak at ∼day 7 (Figure 3C). Senescence is induced in myofibroblasts through the matricellular protein CCN1. During the first three days after becoming senescent, myofibroblasts produce a fibrogenic SASP consisting of PDGF and ECM. Due to this SASP regulation, ECM plateau is still maintained between ∼day 4 to 7, as shown in Figure 3A, despite the drop in myofibroblast numbers. This fibrogenic SASP is also accompanied by juxtacrine-mediated induction of secondary senescence in neighbouring cells, which rises at ∼day 4.5 and begins to drop at ∼day 7.5 (Figure 3K). The model shows that this fibrogenic SASP phase of senescent myofibroblasts helps to maintain the existing fibrogenic state of the proliferative phase while controlling myofibroblast population through the induction of secondary senescence.

Senescent myofibroblasts then shift to the production of a fibrolytic SASP consisting of CSF1, inflammatory SASP and MMP as shown by the sharp rises in these chemical fields at ∼day 7 (Figure 3H-J). The significance of the prior short-lived fibrogenic phase is evident, since with the onset of the fibrolytic phase at ∼day 7, PDGF levels begin to drop 0.6-fold, along with a 0.5-fold drop in ECM levels (Figure 3A and F). The fibrolytic activity of senescent myofibroblasts following ∼day 8 is crucial for wound remodelling, where ECM is produced and broken down eventually coming to an equilibrium as seen with the second plateau in ECM levels (Figure 3A). Inflammatory SASP field also promotes M1 polarisation of wound macrophages, further increasing MMP production to break down scar tissue. This phase is also accompanied by paracrine-mediated induction of senescence, which rises at ∼day 9 just as levels of juxtacrine-induced senescent cell levels begin to decrease. Interestingly, compared to the almost 40-fold increase in juxtacrine-induced senescent cells, paracrine-induced cell number remained rather low with just less than 10-fold increase.

Most senescent cells are eventually removed by immune surveillance mechanisms such as macrophage phagocytosis. Nevertheless, some senescent cells continue to persist long after the wound has healed, where they continue to participate in wound remodelling (Figure 3C). Because of this, MMP levels also remain high to break down excess ECM (Figure 3I), along with inflammatory SASP levels which attract immune cells to further clear senescent cells (Figure 3H). Macrophages eventually undergo apoptosis at the time CSF1 levels drop due to growth factor depletion at ∼day 18, when the wound is sufficiently clear of senescent cells (Figure 3E and J).

### Model classifications reveal that varying spatiotemporal dynamics of senescent cells lead to distinct tissue repair outcomes

To demonstrate the full range of senescent cell function within wound healing, a model of healthy wound healing with baseline parameters was first developed. This model was used to explore the spatiotemporal dynamics of senescent cells within healthy wound healing. Successful wound healing with appropriate levels of ECM, cell composition, and cytokine and growth factor concentrations depends on multiple model parameters. Senescent cell dynamics within the model were primarily controlled by two parameters. These are the probability of senescence in myofibroblasts, which is dictated by the parameter *P_SNC_* and the time of induction of senescent cells in the model, determined by the time constraint parameter *T_SEN_*. Varying these two parameters around their healthy healing simulation baseline values (see Supplementary Table 1) revealed four different scenarios of dysregulated tissue repair ranging from chronic wound to fibrosis. These classifications were based on transient model dynamics and the final simulation state at 18 days.

### Chronic wound inflammation resulting from an increased percentage of senescent cells

The probability of myofibroblast senescence *P_SNC_* was increased from 0.15 to 0.75 to examine the model’s response to a high percentage of senescent cells, with no changes made to the time of senescence induction parameter *T_SEN_*. The increased number of senescent cells, as shown in Figure 5C, resulted in a chronic wound outcome as shown in Figure 4. Although, chronic wounds involve several other dysregulated mechanisms, no other parameters were changed here with the aim of exclusively investigating the effects of the senescent cell population size and dynamics on the repair process. This chronic wound model classification is characterised by insufficient ECM production (Figure 5A), and an accumulation of inflammatory and fibrolytic factors namely, CSF1 (Figure 5I), inflammatory SASP (Figure 5G) and MMP (Figure 5H), compared to its healthy wound healing model counterpart. The corresponding spatial configurations of these characteristics are shown in Figure 4: Senescent cells and macrophages accumulate at the chronic wound site as a result of inefficient immunosurveillance, because the wound macrophages are unable to appropriately handle the high senescent cell load in this simulation. The simulation snapshots also show insufficient ECM deposition; additionally, MMP, inflammatory SASP and CSF1 accumulate at the wound site, whereas total PDGF levels are reduced especially at day 18.

**Figure 4.**
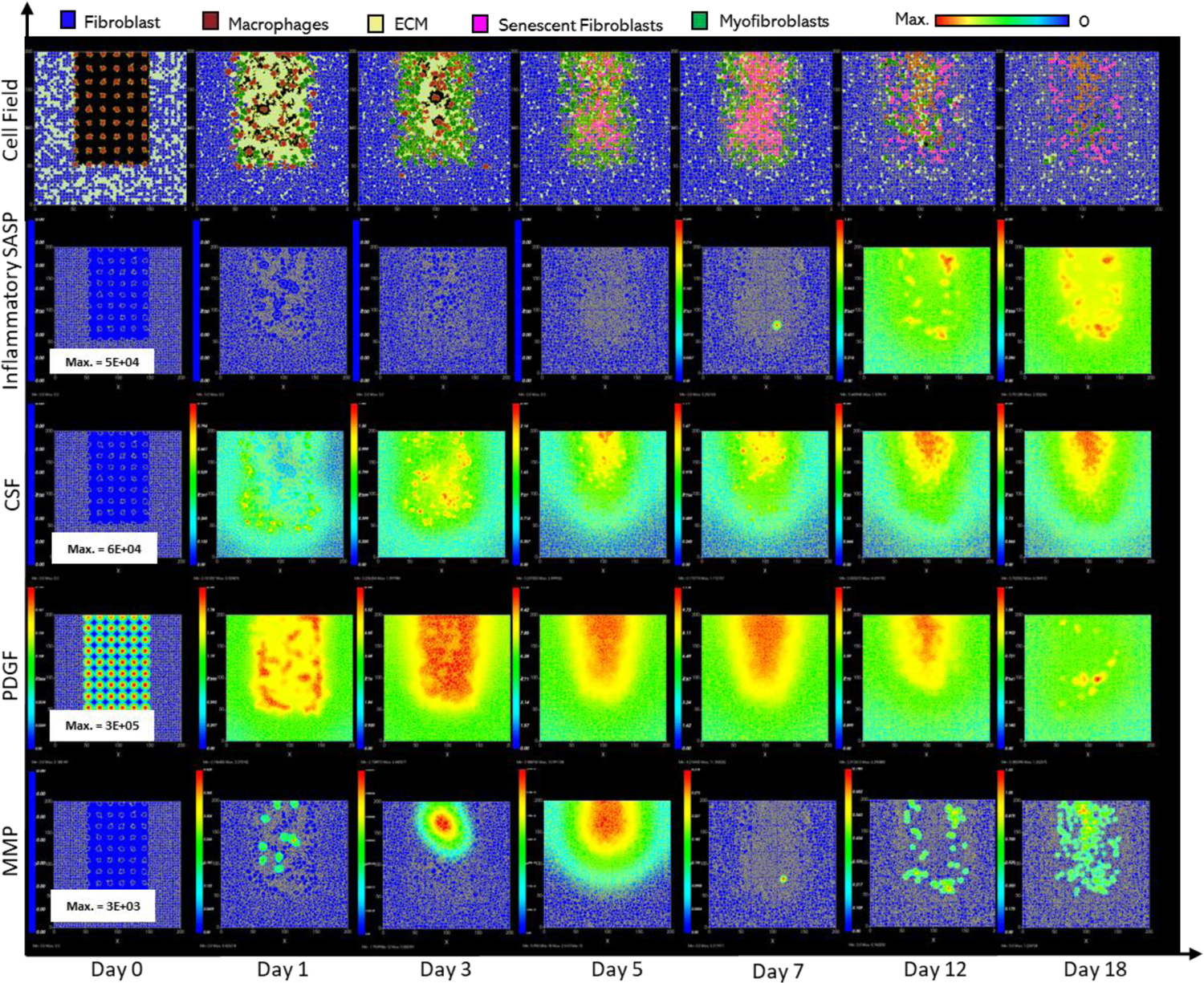
Simulation of chronic wound dynamics in a dermal patch of size 600 µm x 600 µm due to a high percentage of senescent cells. This figure shows snapshots of spatial configuration *vs* time for the progression of a simulated wound due to an excessive number of senescent cells, which results in a chronic wound. The model includes fibroblasts, myofibroblasts, macrophages, ECM and senescent myofibroblasts, and chemical species including inflammatory SASP, CSF1, PDGF and MMP. **Top row:** Snapshots of the simulation cell field at different time points with fibroblasts (blue), myofibroblasts (green), macrophages (brown), ECM (yellow) and senescent myofibroblasts (pink). Snapshots of simulation chemical fields are shown from the second to the last row; **Second row:** Inflammatory SASP, **Third row:** CSF1, **Fourth row:** PDGF, **Fifth row:** MMP. Fields are shaded where, red corresponds to the maximum value specified in the first snapshot panel for each field, and blue corresponds to zero (shown in the colour bar at the top). Snapshots are shown for the time points, left to right, day 0, 1, 3, 5, 7, 12 and 18.

**Figure 5.**
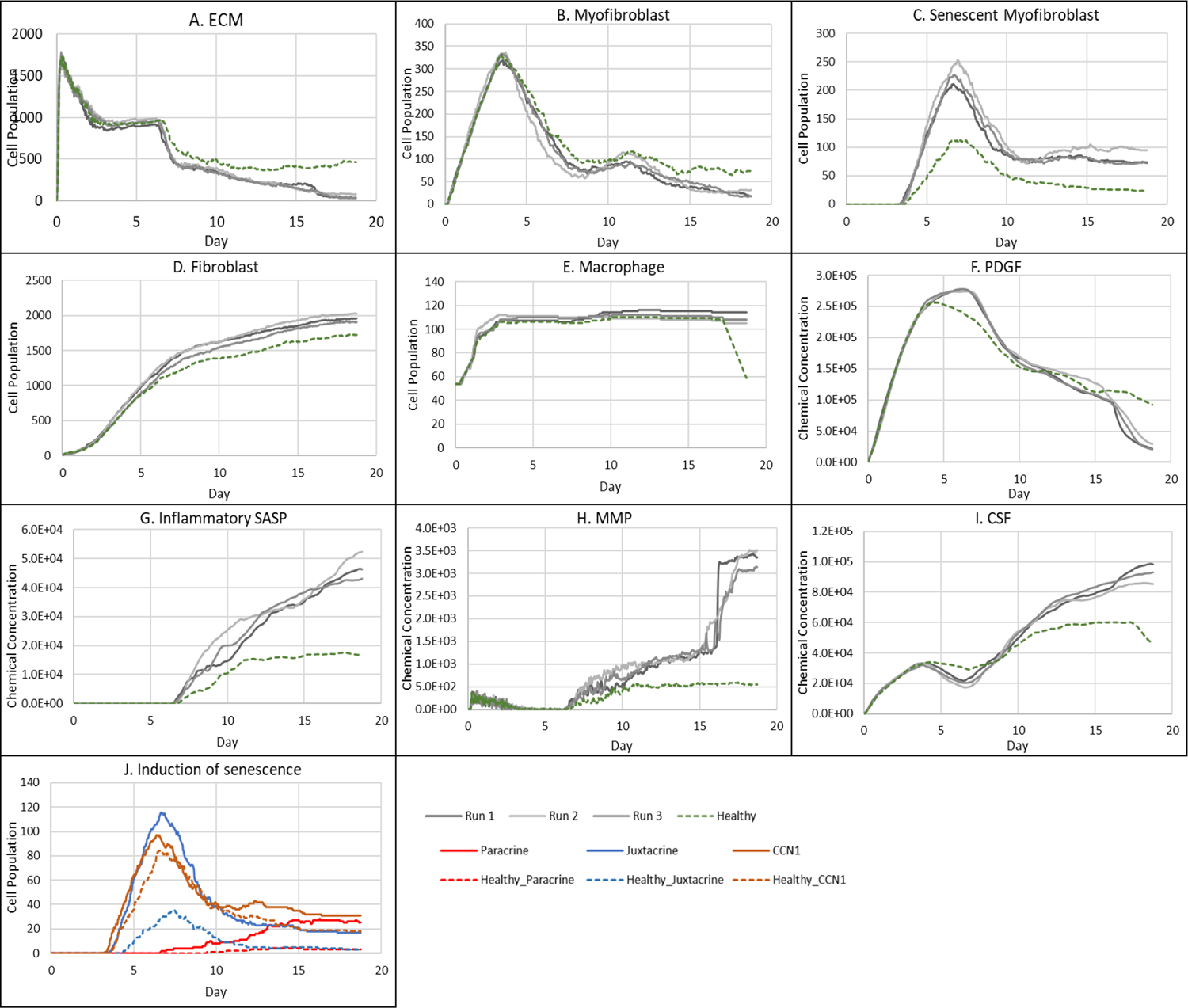
Simulation time series for chronic wound dynamics in a dermal patch of size 600 µm x 600 µm due to a high percentage of senescent cells, shown for model cell types, rate of wound closure and model chemical fields. This figure shows the simulation time series for the progression of a simulated chronic wound due to an excessive number of senescent cells, which results in a chronic wound. Number of cells and total chemical field are shown on the y axis and time (day) is shown on the x axis. Solid lines represent replicate simulation time series for the chronic wound scenario, and the dotted lines in all plots represent simulation time series from the healthy physiological wound healing model (Figure 3), which is included for comparison. Plot are shown for **A:** ECM, **B:** Myofibroblasts, **C:** Senescent myofibroblasts, **D:** Fibroblast, **E:** Macrophages, **F:** PDGF, **G:** Inflammatory SASP, **H:** MMP, **I:** CSF1 and **J:** Induction of senescence. For the time series plots A-I, the three shades of grey represent the different replicate runs as indicated in the legend (since the model is stochastic). For the plot J, different colours represent the different mechanisms of senescence induction, as indicated in the legend; red: paracrine secondary senescence, blue: juxtacrine secondary senescence and orange: CCN1-induced primary senescence. Replicate runs are not shown for the different mechanisms for ease of readability, but total senescent cell numbers for the three replicate runs are shown in plot C.

ECM production during the late healing stages, which is when most matrix remodelling occurs, is significantly lower compared to its healthy counterpart by day 18 (Figure 5A). The decreased amount of ECM can be attributed to increased production of MMP (Figure 5H) by inflammatory M1-polarised macrophages, and senescent cells as shown by the rise in paracrine senescence compared to the healthy healing model (Figure 5J); as discussed previously, paracrine senescence is prevalent during the inflammatory/fibrolytic SASP phase of senescence. This is coupled with attenuated ECM production from fibroblasts and myofibroblasts due to the drop in PDGF around this time (Figure 5F), in line with the decreased PDGF production because of M2 inflammatory macrophage polarisation.

The decreased production of PDGF also leads to reduced myofibroblast proliferation following day ∼11 compared to healthy healing (Figure 5B). However, initial myofibroblast numbers are maintained close to its healthy counterpart since Notch phase (fibrogenic SASP phase) senescent cells promote myofibroblast differentiation through the production of PDGF and matrix components [15, 16]. The following decline occurs as senescent cells shift towards an inflammatory fibrolytic phenotype. Furthermore, macrophage numbers remain elevated, compared to the drop in healthy healing at day ∼17.5 (Figure 5E). This is the consequence of elevated CSF1 levels from SASP, which along with the inflammation and MMP fields, begins to rise around day ∼6 (Figure 5G, H, I).

Inflammatory SASP has been shown to induce paracrine senescence in surrounding cells [32]. Consistently, paracrine senescence is more prevalent in this scenario compared to healthy healing cells (Figure 5J). Interestingly, the number of fibroblasts within the wound region is slightly increased in this classification (Figure 5D).

The inflammatory and proteolytic activity of senescent cells is usually strongly controlled by macrophage immunosurveillance. However, in this classification, the senescent cell burden is too large for macrophage-mediated clearance resulting in impaired healing.

### Fibrotic wound response in the absence of senescent cells

To investigate repair mechanisms in response to a complete removal of senescent cells, the probability of myofibroblast senescence *P_SNC_*was set to zero. The lack of senescent cells (Figure 7C) resulted in a fibrotic response with excessive ECM deposition (Figure 6). The spatial configuration for this response is shown in Figure 6, characterised by: scar tissue formation due to a high level of ECM deposition as shown in the cell field panel; lack of inflammatory SASP due to the absence of senescent cells; high and persistent levels of PDGF; and decreased total MMP levels.

**Figure 6.**
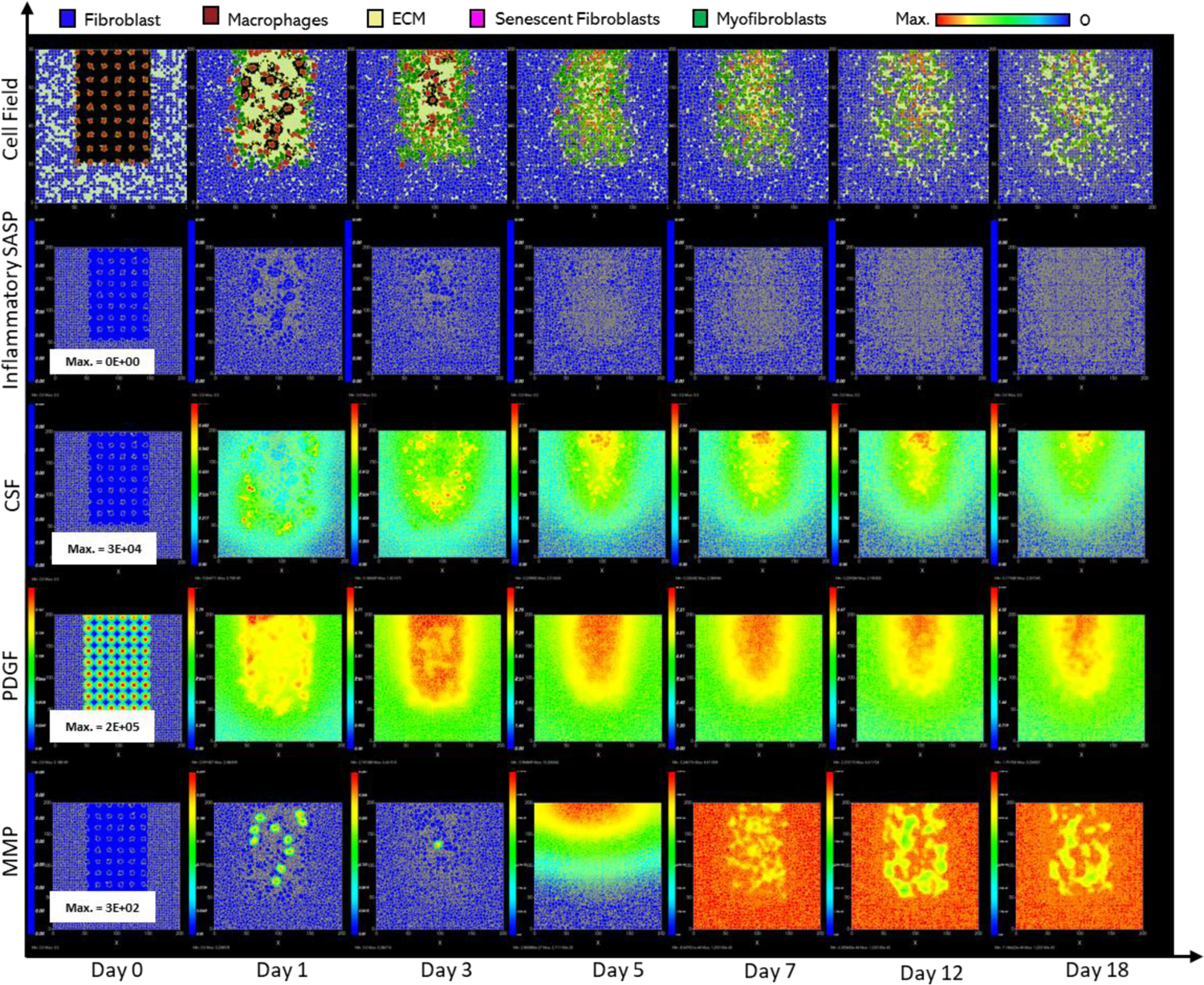
Simulation of fibrotic wound dynamics in a dermal patch of size 600 µm x 600 µm due to the absence of senescent cells. This figure shows snapshots of spatial configuration *vs* time for the progression of a simulated fibrotic wound due to the absence of senescent cells, which results in a fibrotic response. The model includes fibroblasts, myofibroblasts, macrophages and ECM, as well as chemical species including CSF1, PDGF and MMP. **Top row:** Snapshots of the simulation cell field at different time points with fibroblasts (blue), myofibroblasts (green), macrophages (brown) and ECM (yellow). Snapshots of simulation chemical fields are shown from the second to the last row; **Second row:** Inflammatory SASP (which is at zero due to the absence of senescent cells; other types of inflammation were not included in this model), **Third row:** CSF1, **Fourth row:** PDGF, **Fifth row:** MMP. Fields are shaded, where red corresponds to the maximum value specified in the first snapshot panel for each field, and blue corresponds to zero (shown in the colour bar at the top). Snapshots are shown for the time points, left to right, day 0, 1, 3, 5, 7, 12 and 18.

**Figure 7.**
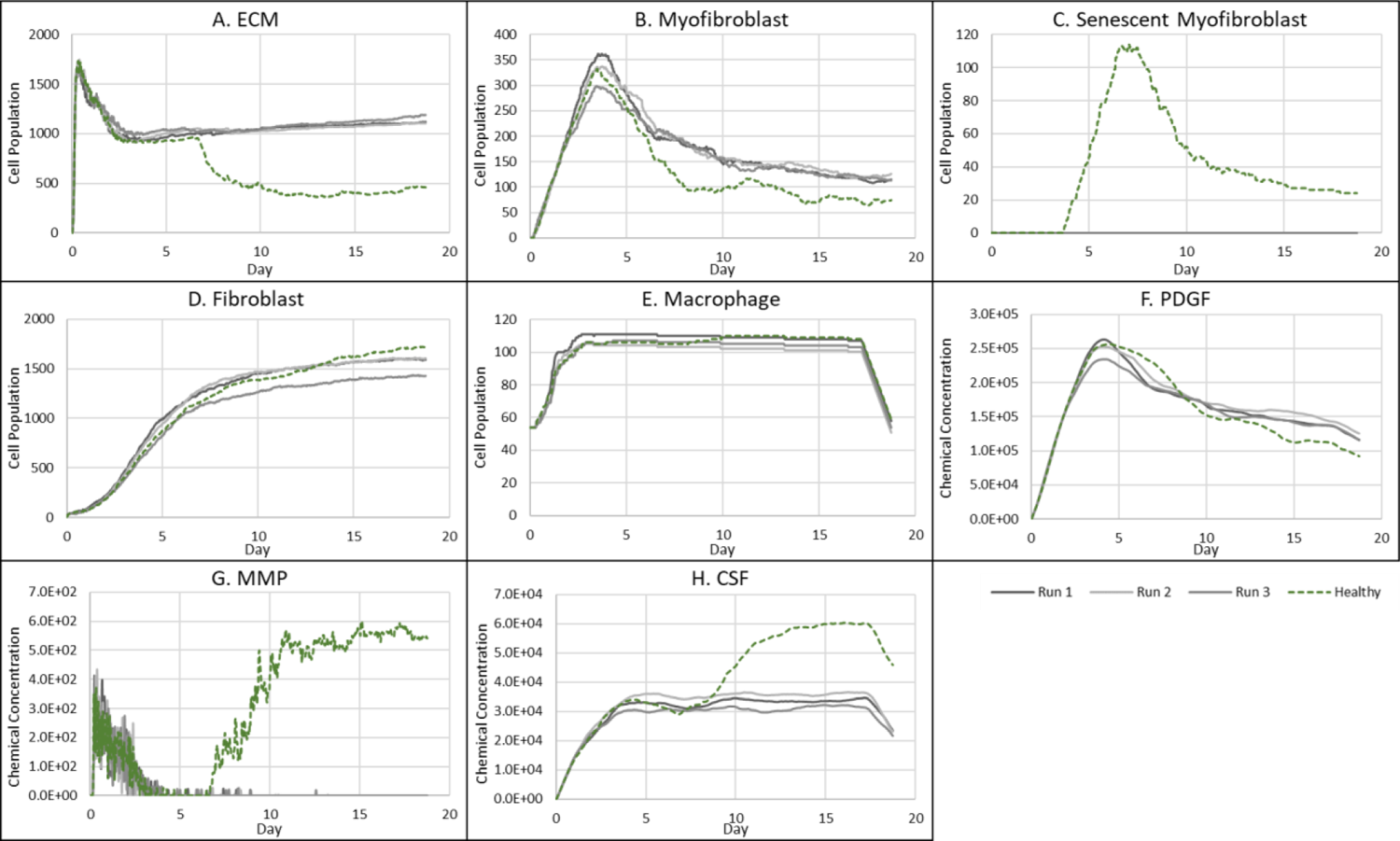
Simulation time series for fibrotic wound dynamics in a dermal patch of size 600 µm x 600 µm due to the absence of senescent cells. This figure shows the simulation time series for the progression of a simulated fibrotic wound due to the absence of senescent cells, which results in a fibrotic response in the wound. Number of cells and total chemical field are shown on the y axis and time (day) is shown on the x axis. Solid lines represent replicate simulation time series for the fibrotic wound scenario, and the dotted lines in all plots represent simulation time series from the healthy physiological wound healing model, included for comparison. Plot are shown for **A:** ECM, **B:** Myofibroblasts, **C:** Senescent myofibroblasts, **D:** Fibroblast, **E:** Macrophages, **F:** PDGF, **G:** MMP and **H:** CSF1. For all the time series plots, the three shades of grey represent the different replicate runs as indicated in the legend (since the model is stochastic).

As stated, this classification shows significantly increased ECM levels compared to healthy healing (Figure 7A). This can be explained in part by elevated PDGF concentration (Figure 7F) and myofibroblast numbers (Figure 7B). Additionally, an important factor contributing to this is the significant decrease in MMP levels compared to healthy healing (Figure 7G). Overall, the simulation shows increased myofibroblast presence and PDGF levels coupled with decreased MMP production to counteract excessive ECM production, all leading to scar tissue production. This can be attributed to insufficient pro-inflammatory polarisation of macrophages by senescent cell factors in the wound, which is necessary for fibrolytic mechanisms during the late stages of healing. This is illustrated by attenuated CSF1 levels (Figure 7H) and a lack of pro-inflammatory SASP (Figure 6), which control macrophage proliferation and pro-inflammatory M1 polarisation, respectively. Furthermore, for this classification, the lack of MMP production by senescent cells contributes to the observed impaired ECM breakdown in the scar tissue. Therefore, the removal of senescent cells disrupts the balance between fibrogenic and fibrolytic mechanisms within the wound, both necessary for a healthy and controlled matrix reconstitution.

### Fibrotic wound response because of delayed induction of senescence

To investigate the effect of a delay in the induction of senescence on wound healing, *T_SEN_* was set to 15 days instead of 12 in the healthy healing model. This means that senescent cells will only appear at the wound site during the later stages of repair. Interestingly, this temporal change resulted in a fibrotic response as shown in Figure 8: This fibrotic response is characterised by an accumulation of ECM as well as senescent cells, as shown in the cell field; the delay in senescent cell induction results in delayed inflammatory SASP production, shown to appear in the day 18 panel.

**Figure 8.**
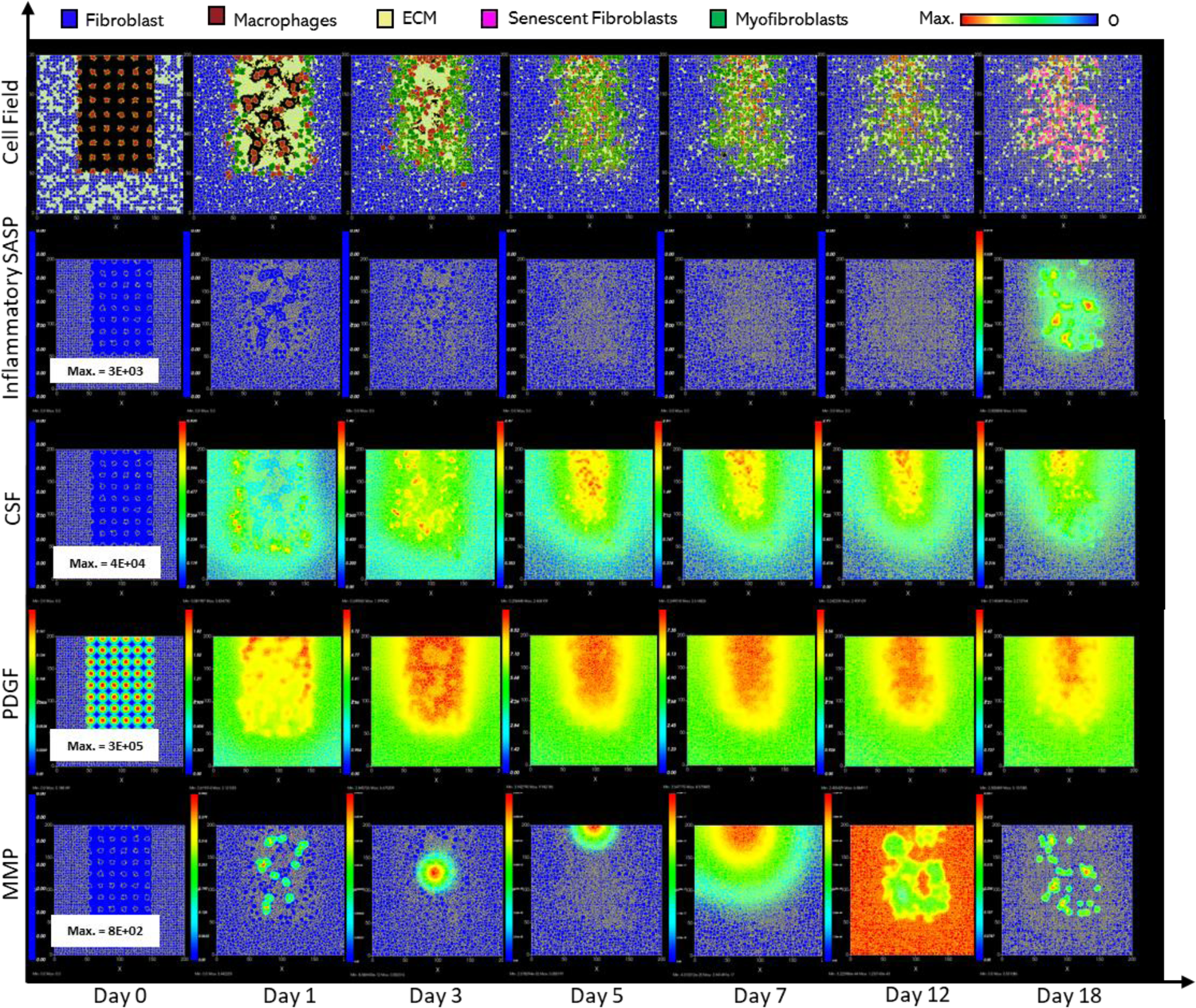
Simulation of fibrotic wound dynamics in a dermal patch of size 600 µm x 600 µm when senescent cell induction is delayed. This figure shows snapshots of spatial configuration *vs* time for the progression of a simulated fibrotic wound due to the delayed induction of senescent cells during the later stages of healing, which results in a fibrotic response. The model includes fibroblasts, myofibroblasts, macrophages, ECM and senescent myofibroblasts, and chemical species including inflammatory SASP, CSF1, PDGF and MMP. **Top row:** Snapshots of the simulation cell field at different time points with fibroblasts (blue), myofibroblasts (green), macrophages (brown), ECM (yellow) and senescent myofibroblasts (pink). Snapshots of simulation chemical fields are shown from the second to the last row; **Second row:** Inflammatory SASP, **Third row:** CSF1, **Fourth row:** PDGF, **Fifth row:** MMP. Fields are shaded, where red corresponds to the maximum value specified in the first snapshot panel for each field, and blue corresponds to zero (shown in the colour bar at the top). Snapshots are shown for the time points, left to right, day 0, 1, 3, 5, 7, 12 and 18.

As expected, senescent cells in this model classification rise around day ∼15.5 (Figure 9C). The resulting fibrotic response is accompanied by elevated ECM levels compared to the healthy healing simulation (Figure 9A). A small drop in ECM is observed, coinciding with an increase in MMP levels at around ∼day 18 (Figure 9H), which otherwise remained significantly low throughout the simulation. This response, along with small increases in inflammatory SASP and CSF1 (Figure 9G-H and inflammatory SASP panel shown in Figure 8) implies the shift of senescent cells to an inflammatory phenotype at this time point. MMP levels peak at day 18 as they are produced by senescent cells in their inflammatory phase and M1 polarised macrophages, which is triggered by this short increase in inflammatory SASP (Figure 9G). However, this fibrolytic activity is too late to attenuate the fibrotic response by breaking down the scar tissue. Furthermore, due to the delay in MMP activity, ECM levels remain elevated despite a drop in myofibroblast levels due to senescence induction (Figure 9B). ECM production here is supported by PDGF levels which remain elevated compared to its healthy counterpart as it is produced by fibrogenic phase senescent cells (Figure 9F). CSF1 levels also remain low (Figure 9I) but do not have a major impact of macrophage numbers (Figure 9E). Aside from CCN1-induced senescent cells, juxtacrine secondary senescence is prevalent (Figure 9J). As discussed, this is in accordance with previous work showing that juxtacrine senescence is accompanied by NOTCH-induced fibrogenic SASP rich in PDGF and ECM components [71]. The lack of paracrine secondary senescent cells here is congruent with the lack of fibrolytic activity in the wound model.

**Figure 9.**
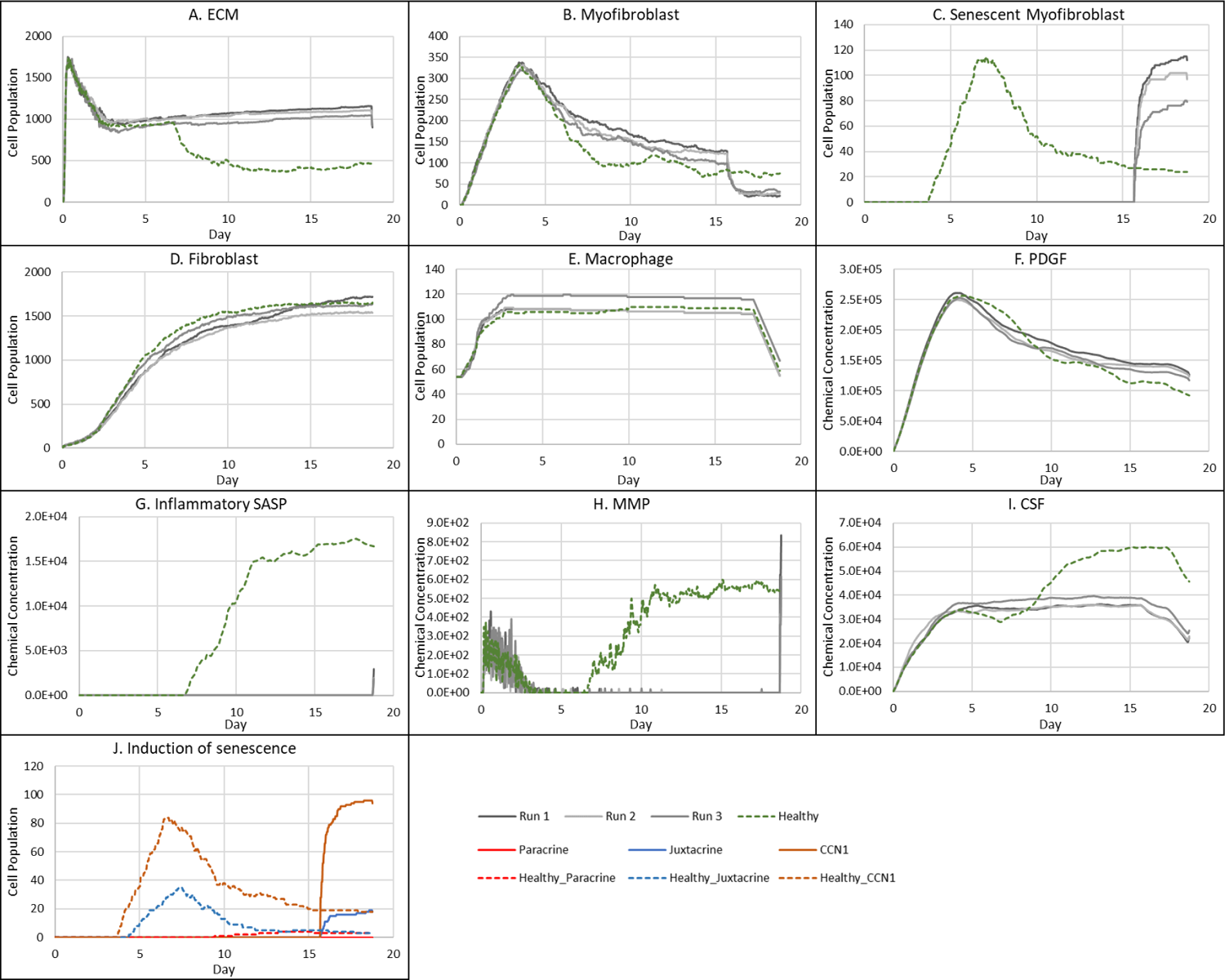
Simulation time series for fibrotic wound dynamics in a dermal patch of size 600 µm x 600 µm when senescent cell induction is delayed. This figure shows the simulation time series for the progression of a simulated fibrotic wound due to the delayed temporal induction of senescent cells during the later stages of healing, which results in a fibrotic response. Number of cells and total chemical field are shown on the y axis and time (day) is shown on the x axis. Solid lines represent simulation time series for this fibrotic wound scenario, and the dotted lines in all plots represent simulation time series from the healthy physiological wound healing model, included for comparison. Plot are shown for **A:** ECM, **B:** Myofibroblasts, **C:** Senescent myofibroblasts, **D:** Fibroblasts, **E:** Macrophages, **F:** PDGF, **G:** Inflammatory SASP, **H:** MMP, **I:** CSF1 and **J:** Induction of senescence. For the time series plots A-I, the three shades of grey represent the different replicate runs as indicated in the legend (since the model is stochastic). For the plot J, different colours represent the different mechanisms of senescence induction, as indicated in the legend; red: paracrine secondary senescence, blue: juxtacrine secondary senescence and orange: CCN1-induced primary senescence. Replicate runs are not shown for the different mechanisms for ease of readability, but total senescent cell numbers for the three replicate runs are shown in plot C.

In this scenario, due to the timing of senescence induction in the wound, the existing fibrogenic activity of the repair process is augmented by senescent cells which are still in their NOTCH-induced fibrogenic SASP phase, rich in PDGF and ECM in the model [32]. This overlap of fibrogenic activity, with a lack of fibrolytic activity leads to an overall fibrotic response. This lack of fibrolytic activity is due to a delay in the shift to an inflammatory senescence phenotype and insufficient macrophage M1 polarisation during late-stage healing, both of which are required to assist in ECM remodelling.

These results show that aside from SASP composition and secondary senescence mechanisms, the temporal aspect of senescence induction is also a crucial factor in repair outcomes. To summarise, the existing fibrogenic wound activity in this scenario was strengthened by senescent cells, which eventually shift to a fibrolytic phenotype. However, the shift is too late to repair the scar tissue due to a disruption in the sequence of repair mechanisms.

Inflammatory SASP from pre-existing senescent cells leads to chronic wound inflammation Next, the added effect of inflammatory SASP from pre-existing senescent cells in wound healing was investigated. This scenario represents senescence induction during the early inflammatory phase of the wound healing process, as well as pre-existing senescent cells as seen with age-related accumulation of senescence. To this end, inflammatory senescent cells were introduced at the start of the model simulation with no other parameter changes (Figure 10). This classification simplistically represents an ageing or a diabetic wound, both of which are accompanied by an accumulation of inflammatory senescent cells, which is shown in the first simulation cell field snapshot at day 0 in Figure 10. High levels of inflammatory SASP are also prevalent at day 0, representing increased systemic inflammation, as seen with age. Similarly, MMP and CSF1 levels are also high throughout, whereas PDGF levels are significantly reduced (Figure 10).

**Figure 10.**
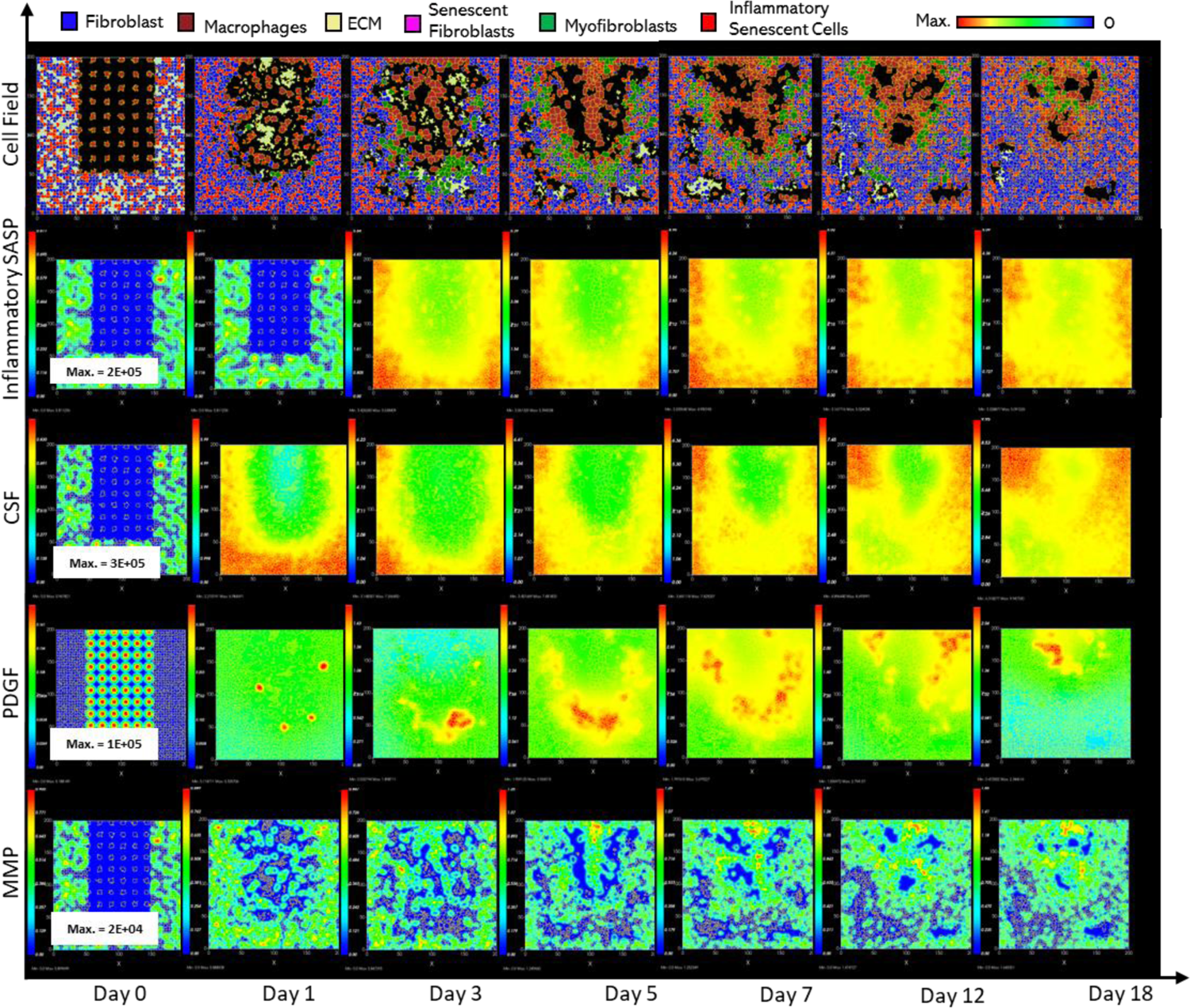
Simulation of chronic wound dynamics in a dermal patch of size 600 µm x 600 µm due to pre-existing (inflammatory) senescent cells. This figure shows snapshots of spatial configuration *vs* time for the progression of a simulated chronic wound due to pre-existing senescent cells which produce inflammatory SASP; this represents wound healing during ageing, which is accompanied by an accumulation of senescent cells, as well as senescence induced during the inflammatory phase of the wound healing process, resulting in a chronic wound. The model includes fibroblasts, myofibroblasts, macrophages, ECM, pre-existing (inflammatory) senescent cells and senescent myofibroblasts, and chemical species including PDGF, MMP, inflammatory SASP and CSF1. **Top row:** Snapshots of the simulation cell field at different time points with fibroblasts (blue), myofibroblasts (green), macrophages (brown), ECM (yellow), pre-existing inflammatory senescent cells (red) and senescent myofibroblasts (pink). Snapshots of simulation chemical fields are shown from the second to the last row; **Second row:** Inflammatory SASP, **Third row:** CSF1, **Fourth row:** PDGF, **Fifth row:** MMP. Fields are shaded, where red corresponds to the maximum value specified in the first snapshot panel for each field, and blue corresponds to zero (shown in the colour bar at the top). Snapshots are shown for the time points, left to right, day 0, 1, 3, 5, 7, 12 and 18.

Senescent cell numbers remained steadily elevated throughout the model simulation after an initial drop in numbers at around ∼day 2, due to macrophage-mediated clearance (Figure 11C). Macrophage immunosurveillance here is mediated by inflammatory SASP produced by the pre-existing senescent cells, which attracts macrophages towards senescent cells in the model. Inflammatory SASP rises sharply at around ∼day 1 followed by a decrease at ∼day 2.5 and then continues at steady levels, with levels much higher compared to the healthy healing simulation (Figure 11G).

**Figure 11.**
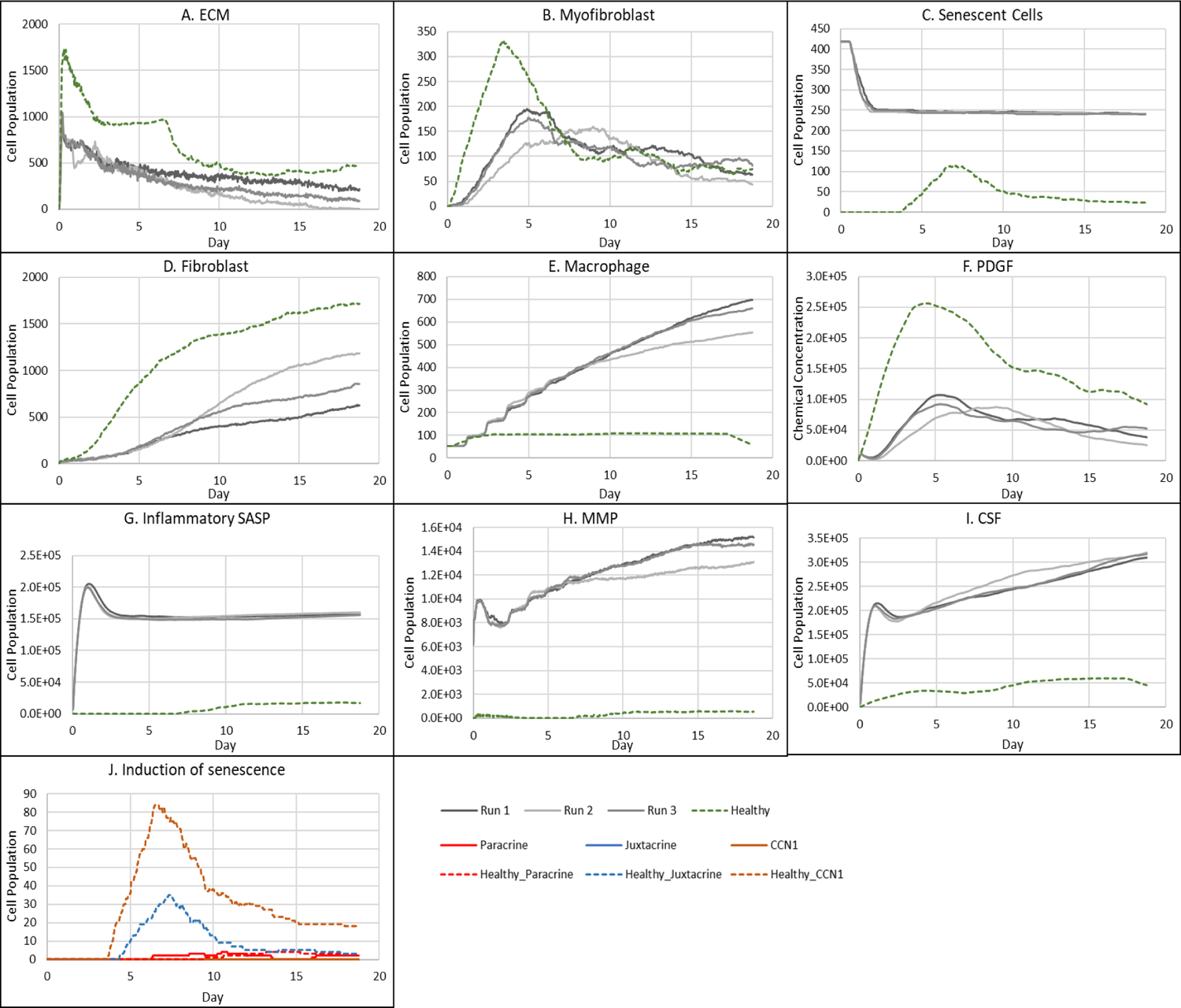
Simulation time series for chronic wound dynamics in a dermal patch of size 600 µm x 600 µm due to pre-existing (inflammatory) senescent cells. This figure shows the simulation time series for the progression of a simulated chronic wound due to pre-existing senescent cells; this represents wound healing during ageing, which is accompanied by an accumulation of senescent cells, as well as senescence induced during the inflammatory phase of the wound healing process, resulting in a chronic wound. Number of cells and total chemical field are shown on the y axis and time (day) is shown on the x axis. Solid lines represent simulation time series for this chronic wound scenario, and the dotted lines in all plots represent simulation time series from the healthy physiological wound healing model, included for comparison. Plot are shown for **A:** ECM, **B:** Myofibroblasts, **C:** Total senescent cells, **D:** Fibroblasts, **E:** Macrophages, **F:** PDGF, **G:** Inflammatory SASP, **H:** MMP, **I:** CSF1 and **J:** Induction of senescence. For the time series plots A-I, the three shades of grey represent the different replicate runs as indicated in the legend (since the model is stochastic). For the plot J, different colours represent the different mechanisms of senescence induction, as indicated in the legend; red: paracrine secondary senescence, blue: juxtacrine secondary senescence and orange: CCN1-induced primary senescence. Replicate runs are not shown for the different mechanisms for ease of readability, but total senescent cell numbers for the three replicate runs are shown in plot C.

The same pattern is observed with other factors produced by senescent cells including MMPs (Figure 11H) and CSF1 (Figure 11I), coinciding with the senescent cell levels. MMP levels remain chronically elevated resulting in excessive breakdown of ECM (Figure 11A). Rising CSF1 levels result in uncontrolled macrophage proliferation (Figure 11E-I). Aside from triggering macrophage immunosurveillance, pro-inflammatory SASP also induces pro-inflammatory macrophage polarisation (Figure 11G). The rising number of inflammatory macrophages also contributes to increased MMP production alongside that of senescent cells.

Pro-inflammatory M1 polarisation of macrophages also means decreased PDGF production, as shown in Figure 11F. This results in decreased fibroblast proliferation (Figure 11D) and overall ECM production by fibroblasts and myofibroblasts (Figure 11A). Myofibroblast population was significantly reduced compared to healthy healing as shown during the first 7 days of the mid healing stage (Figure 11B). This is due to decreased PDGF production by M1 polarised macrophages and inflammatory SASP-producing senescent cells, resulting in decreased myofibroblast differentiation. However, myofibroblast numbers during the later time points match that of healthy healing during which time myofibroblasts usually begin to be cleared out from the wound site.

Insufficient PDGF production leads to attenuated fibroblast levels, as stated previously (Figure 11D). Interestingly, in this classification, the number of fibroblasts was decreased compared to the chronic wound dynamics shown in Figure 5, which could imply decreased fibroblast activation from quiescence. Furthermore, the more prevalent mechanism for secondary senescence in this classification is paracrine, with minimal juxtacrine induction (Figure 11J). This makes sense since paracrine senescence is more associated with an inflammatory SASP compared to juxtacrine senescence, as discussed previously. Lastly, this model classification exhibited greater run-to-run variability (Figure 11).

### Sensitivity analysis reveals that deviation from the tightly controlled senescence program during wound healing leads to dysregulated repair mechanisms

To explore any trade-offs between the population size of senescent cells and the time of senescence induction during wound healing, a multidimensional parameter sweep of probability of myofibroblast senescence *P_SNC_*and senescence induction time constraint *T_SEN_* was performed (see methods). Simulations were run for each parameter set by increasing and decreasing parameter values around their baseline values. For each simulation, cell population and chemical concentrations were examined, as shown for ECM in Figure 13, and for myofibroblasts, macrophages, senescent cells, PDGF, CSF1, Inflammatory SASP and MMP in Supplementary Figure 2 - Supplementary Figure 2. Three regions of the parameter space were identified as shown by the final simulation states at day 18 of the parameter sweep in Figure 12, where blue indicates fibrotic wound, red indicates chronic inflammation and green indicates healthy healing.

**Figure 12.**
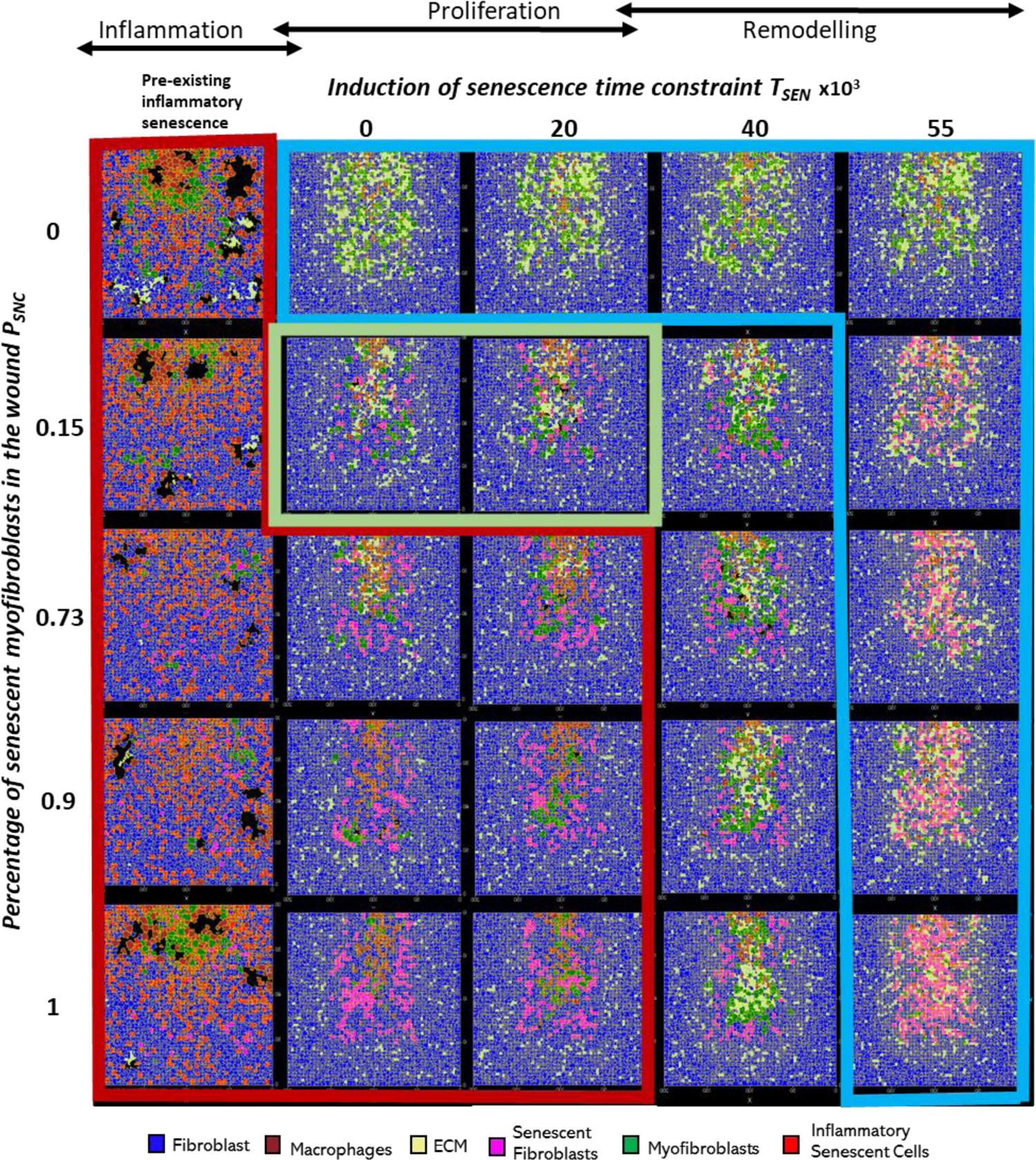
Wound healing model simulation snapshots of the last time point from multidimensional parameter sensitivity analysis for variations in the parameters probability of myofibroblast senescence (*P_SNC_*) and senescence induction time constraint (*T_SEN_*) showing regions with distinct repair dynamics. This plot shows the model simulation snapshots of the final time point ∼day 18 from a pairwise parameter sweep of the parameters: percentage of myofibroblast senescence *P_SNC_* (shown vertically on the left) and senescence induction time constraint *T_SEN_* (shown horizontally at the top), around their baseline values (provided in Supplementary Table 1). Along with the different values included in the parameter sweep for the senescence induction time constraint *T_SEN_* parameter, pre-existing inflammatory senescent cells are also shown to represent senescence induced during the inflammatory phase of the wound healing process (i.e., before *T_SEN_* = 0) which was not explicitly included in this model. Arrows above the *T_SEN_* parameter sweep values show the wound healing phases (inflammation, proliferation and remodelling phases) represented by the *T_SEN_* parameter values. The cell types in the snapshots are fibroblasts (blue), myofibroblasts (green), macrophages (brown), ECM (yellow), pre-existing (inflammatory) senescent cells (red) and senescent myofibroblasts (pink). The boxes highlight regions with distinct repair dynamics for chronic wound inflammation (red border box), healthy healing (green border box) and fibrotic wound response (blue border box).

**Figure 13.**
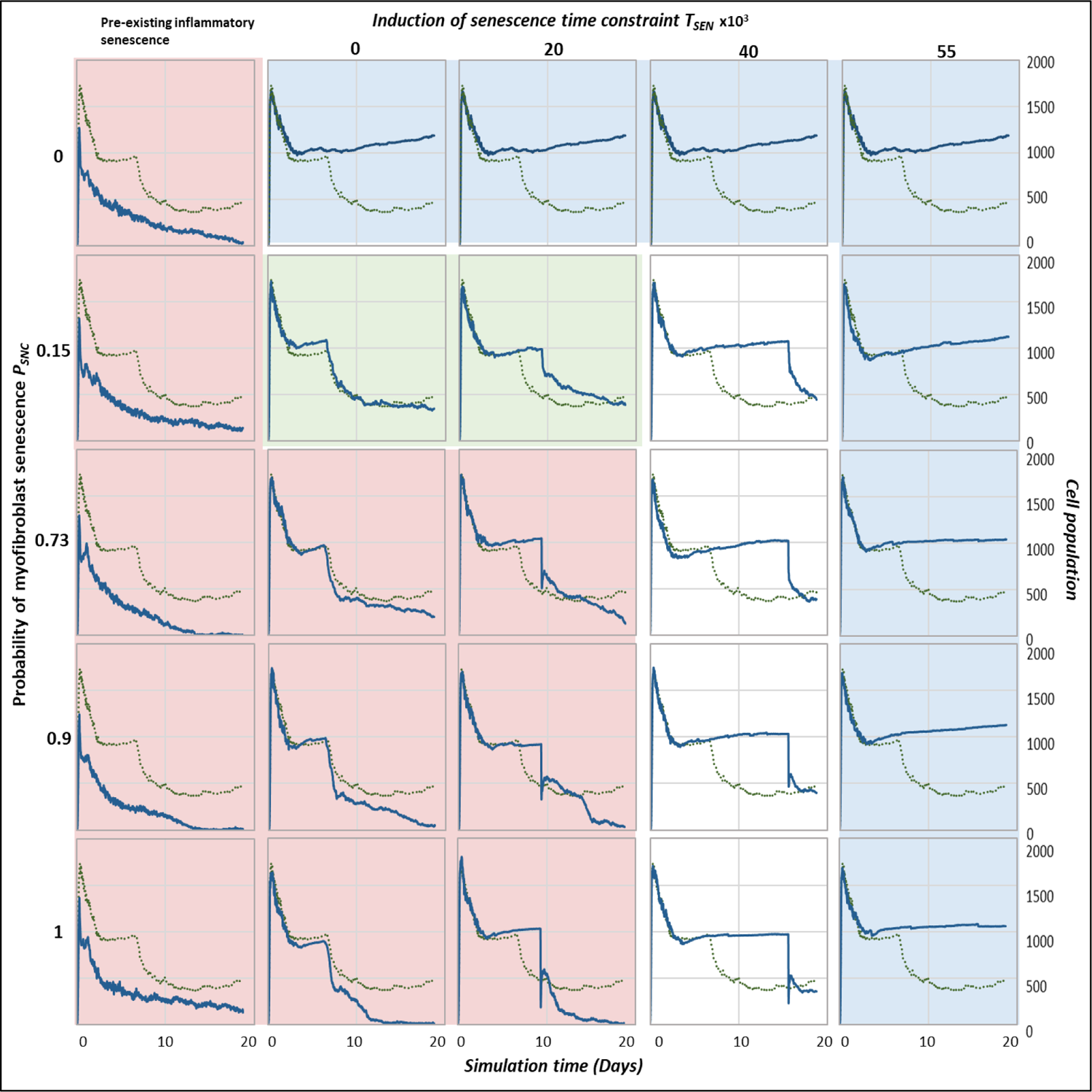
Wound healing model multidimensional sensitivity analysis of the amount of ECM vs time for variations in the parameters probability of myofibroblast senescence (*P_SNC_*) and senescence induction time constraint (*T_SEN_*) showing regions with distinct repair dynamics. The subplots show the model simulation time series for ECM levels in the wound from a pairwise parameter sweep of the parameters: probability of myofibroblast senescence *P_SNC_* (shown vertically on the left) and senescence induction time constraint *T_SEN_* (shown horizontally at the top), around their baseline values (provided in Supplementary Table 1). Along with the different values included in the parameter sweep for the senescence induction time constraint *T_SEN_* parameter, pre-existing inflammatory senescent cells are also shown to represent senescence induced during the inflammatory phase of the wound healing process (i.e., before *T_SEN_* = 0) which was not explicitly included in the model. Simulation time in days is shown on the x axis and cell population numbers are shown on the y axis. The boxes highlight regions with distinct repair dynamics: chronic wound inflammation (red shaded box), healthy healing (green shaded box) and fibrotic wound response (blue shaded box). The solid blue line represents simulation time series from the pairwise parameter sweep. The dotted lines in all the plots represent simulation time series from the healthy physiological wound healing model were included for comparison.

For all values considered for the probability of myofibroblast senescence *P_SNC_*, the presence of pre-existing senescent cells in the wound site (or the induction of senescence during the inflammatory phase of healing) leads to chronic inflammation and an inability for the wound to resolve (red boxed and shaded subplots in Figure 12, Figure 13 and Supplementary Figure 2-Supplementary Figure 8). As shown in the plots, the chronic wound formation is due to insufficient ECM production, myofibroblast differentiation and PDGF levels, along with high levels of macrophages, MMP, CSF1 and inflammatory SASP. Most cellular and chemical components are similar except for the mechanism of secondary senescence, where higher *P_SNC_* values are accompanied by higher numbers of paracrine-mediated secondary senescence. Overall, pre-existing inflammatory senescence is accompanied by increased paracrine senescence compared to juxtacrine. This may be expected since SASP in chronic wound is rich in inflammatory and proteolytic factors but deficient in growth factors and matrix proteins [28, 72, 73].

When *P_SNC_* was set to zero (blue boxed and shaded subplots top row in Figure 12, Figure 13 and Supplementary Figure 2-Supplementary Figure 8), the simulation resulted in a fibrotic response characterised by excessive ECM deposition and lack of the fibrolytic activity that would usually be mediated by senescent cells and inflammatory M1 polarised macrophages. On the other hand, simulations run with low *P_SNC_* and low *T_SEN_* values result in healthy healing (green boxed and shaded subplots in Figure 12, Figure 13 and Supplementary Figure 2-Supplementary Figure 8). The cell populations and chemical concentrations in these simulations are consistent with their healthy healing counterparts. However, ECM levels remain elevated for a longer duration, until day ∼9, for the simulation with a low to moderate *T_SEN_* value compared to a low *T_SEN_* value. This is because the short delay in senescence induction leads to overall ECM production in the wound being augmented by senescent cell-derived ECM and PDGF. This does not lead to a fibrotic state since senescent cell and macrophage derived MMP is able to attenuate ECM levels sufficiently and on time.

In contrast, for high *P_SNC_* and high *T_SEN_* values (blue boxed and shaded subplots bottom right in Figure 12, Figure 13 and Supplementary Figure 2-Supplementary Figure 8), simulations result in fibrotic states, due to an increased number of senescent myofibroblasts that are induced during the late stages of healing. Myofibroblast numbers in these simulations remain significantly elevated compared to healthy healing, as do ECM levels. This is because the fibrogenic SASP phase of senescent cells is augmenting the intrinsic fibrogenic activity of the wound. This happens through sustained anti-inflammatory macrophage activity whereby they continue to produce PDGF, which is also elevated, for fibroblasts and myofibroblasts. A lack of macrophage pro-inflammatory polarisation leads to insufficient MMP production. Myofibroblasts and fibroblasts continue to produce excessive amounts of ECM in an uncontrolled manner resulting in the scar tissue as shown. This behaviour is consistently observed regardless of the myofibroblast senescence probability parameter *P_SNC_* (blue boxed and shaded subplots top right in Figure 12, Figure 13 Supplementary Figure 2-Supplementary Figure 8). That is, simulations with low *P_SNC_* and high *T_SEN_* values also result in a fibrotic wound state. This highlights the importance of time of induction of senescent cells during the repair process. In all these simulations, juxtacrine induction is the dominant mode of secondary senescence, which is to be expected given that fibrogenic SASP is associated with juxtacrine senescence, as discussed previously several times.

On the other hand, for the moderate to high *T_SEN_* value and low *P_SNC_* value (topmost white subplots in Figure 12, Figure 13 and Supplementary Figure 2-Supplementary Figure 8), the fibrotic response is elevated until day ∼15, after which it is rescued by the switch to a fibrolytic SASP in the senescent cells, which is indicated by the sharp rise in inflammatory SASP, MMP and CSF1, and a drop in PDGF levels. This is also accompanied by macrophages switching to a pro-inflammatory phenotype, contributing to the drop in PDGF and rise in MMP. However, for the same *T_SEN_* value but high *P_SNC_* values (second to last white subplots in Figure 12, Figure 13 and Supplementary Figure 2-Supplementary Figure 8), simulations are slightly more fibrotic. These are accompanied by sustained myofibroblast activity and higher inflammatory SASP compared to healthy healing. However, these also eventually resolve.

Lastly, simulations run with high *P_SNC_* and low *T_SEN_* values consistently result in a chronic wound state (red boxed and shaded subplots in Figure 12, Figure 13 and Supplementary Figure 2-Supplementary Figure 8). The end of these simulations is accompanied by decreased ECM production and myofibroblast numbers overall. Macrophage numbers remain elevated towards the end of the simulations, when they usually decrease in healthy healing. This is due to increased CSF1 production by inflammatory senescent cells. Increased inflammation mediated by senescent cells is also characterised by higher inflammatory SASP levels, increased MMP and decreased PDGF levels. Moreover, paracrine secondary senescence is also more prevalent compared to healthy healing. Altogether, increased ECM breakdown and accumulation of inflammatory factors result in an improper wound microenvironment and tissue composition. This is because of a high senescent cell burden that is unable to be resolved by macrophage-mediated immunosurveillance despite their pro-inflammatory polarisation.

## Discussion

Despite their crucial role in physiological tissue repair and in mitigating fibrotic disorders, senescent cells are also involved in the pathogenesis of fibrotic diseases [22–24] and chronic wounds[25–27]. Senescent cells play a heterogenous and sometimes paradoxical role in tissue repair. This could be due to the different effects of the duration of the senescence program; transient senescence is beneficial, conferring pro-healing factors, whereas prolonged senescence is detrimental, leading to dysregulated immunosurveillance mechanisms and disrupted tissue repair [74]. The wound healing model developed here was used here to show that aside from this duration of senescence, other factors including dynamic SASP composition and timing of senescence induction, relative to the repair phase, are also important to consider as they steer healing dynamics towards different outcomes.

To demonstrate the full range of senescent cell functions within dysregulated tissue repair, variations in senescent cell spatiotemporal dynamics were investigated. Senescent cell dynamics within the model are primarily controlled by two parameters. These are the probability of senescence in myofibroblasts, which is dictated by the parameter *P_SNC_*, and the time of induction of senescent cells in the model, determined by the time constraint parameter *T_SEN_*. Varying these two parameters around their healthy healing simulation baseline values revealed four distinct classifications of dysregulated tissue repair due to senescent cell spatiotemporal dynamics ranging from chronic wound to fibrosis. To summarise, an increased percentage of background senescent cells in the wound microenvironment results in a chronic wound, whereas the absence of senescent cells results in a fibrotic wound. Interestingly, the delayed induction of senescent cells during the late stages of healing also results in a fibrotic wound. Lastly, the early induction of senescent cells during the inflammation phase or the presence of systemic pre-existing inflammatory senescence, which represents prolonged senescence in an aged tissue for instance, results in a chronic wound.

A multidimensional parameter sensitivity analysis showed how the interaction between the probability of myofibroblast senescence and time of senescence induction can lead to different outcomes. Simulations with pre-existing pro-inflammatory senescence for all values of probability of myofibroblast senescence (*P_SNC_*) resulted in chronic inflammation (Figure 12, Figure 13 and Supplementary Figure 2-Supplementary Figure 8 red shaded subplots). The features of chronic wound dynamics in these simulations confirmed decreased fibrogenic activity, characterised by insufficient ECM production, myofibroblast differentiation and PDGF levels, as previously reported in chronic wounds [75, 76]. Additionally, increased proteolytic and inflammatory activity was apparent in these simulations, namely, increased number of macrophages, and high wound concentrations of MMP, CSF1 and inflammatory SASP, in line with previous observations in clinical studies of chronic wounds [77–79]. Intriguingly, pre-existing inflammatory senescence is accompanied by increased paracrine secondary senescence compared to juxtacrine senescence, with the difference augmented with increasing numbers of senescent cells. This result within the model simulations may be expected since paracrine senescence is associated with SASP rich in inflammatory and proteolytic components [28, 72, 73, 80].

Similarly, simulations with high numbers of senescent cells induced during the mid stages of healing (high *P_SNC_* and low *T_SEN_* values) also consistently result in a chronic wound state, which has been observed in chronic wound mouse models [28] (Figure 12, Figure 13 and Supplementary Figure 2-Supplementary Figure 8 red shaded subplots). These scenarios were characterised by decreased ECM production and myofibroblast numbers, and elevated inflammatory and proteolytic factors. These simulations were also accompanied by increased paracrine secondary senescence, compared to healthy healing. With increasing *P_SNC_* values, paracrine senescence dominated over juxtacrine secondary senescence. The senescent cell burden in these simulations was too overpowering to be resolved by macrophage-mediated immunosurveillance. Chronic wound inflammation due to a higher percentage of senescent cells, interestingly, exhibited enhanced fibroblast proliferation, compared to the chronic wound resulting from pre-existing inflammatory senescent cells which represents an aged tissue.

Contrastingly, the outcome of simulations with no senescent myofibroblasts (*P_SNC_* is set to zero, excluding pre-existing inflammatory senescent cells) was a fibrotic response characterised by excessive ECM deposition and lack of fibrolytic activity, which is typically observed in fibrosis [62, 81–83] (Figure 12, Figure 13 and Supplementary Figure 2-Supplementary Figure 8 blue shaded subplots). This agrees with the fact that, aside from being a source of proteolytic enzymes during repair, senescent cells also produce factors that promote pro-inflammatory macrophage polarisation [44]. The polarisation of macrophages to a M1 phenotype is important during the remodelling stages since it selectively upregulates several MMPs involved in breaking down excessive scar tissue during healing [84].

A similar fibrotic response is observed in simulations with a high number of senescent cells induced during the late stages of repair (high *P_SNC_* and high *T_SEN_* values) (Figure 12, Figure 13 and Supplementary Figure 2-Supplementary Figure 8 blue shaded subplots bottom right). This response is also characterised by typical fibrosis features including elevated myofibroblast numbers and ECM levels. In these simulations the fibrogenic SASP phase of senescent cells is adding to the fibrogenic activity of the repair process. This takes place through sustained anti-inflammatory macrophage activity, whereby they continue to produce PDGF, which is also elevated for fibroblasts and myofibroblasts. A lack of macrophage pro-inflammatory polarisation leads to insufficient MMP production. Myofibroblasts and fibroblasts continue to produce excessive amounts of ECM in an uncontrolled manner resulting in the scar tissue formation. Interestingly, this behaviour is seen consistently regardless of the myofibroblast senescence probability parameter *P_SNC_*. This is shown in simulations with the appropriate initial amount of senescent cells but time of senescence induction is during the late repair stage (low *P_SNC_*with high *T_SEN_* values) (Figure 12, Figure 13 and Supplementary Figure 2-Supplementary Figure 8 blue shaded subplots top right). This highlights the importance of timing in the induction of senescent cells during the repair process relative to the repair stage, aside from SASP composition itself and the total duration of senescence. In all these simulations, juxtacrine induction is the dominant mode of secondary senescence, which is to be expected given that fibrogenic SASP is associated with juxtacrine senescence.

Lastly, simulations run with a small number of senescent cells during the mid stages of repair (low *P_SNC_*and low *T_SEN_* values) result in healthy wound healing (Figure 12, Figure 13 and Supplementary Figure 2-Supplementary Figure 8 green shaded subplots). In these simulations, the presence of transient senescence during the mid stages of healing represented by low *T_SEN_* values, promotes healing as shown by the simulation time series data.

Senescent cells initially promote myofibroblast differentiation and ECM production, but then shift to promote excess ECM degradation, upregulation of immune surveillance through pro-inflammatory SASP and M1 macrophage polarisation [30–32, 44]. These processes take place in a timely manner with respect to the repair process in place, resulting in healthy healing. Even a slight deviation is shown to result in some scarring, as observed with the simulation with the higher *T_SEN_* value. This is because ECM production in the wound is augmented by senescent cell-derived ECM and PDGF. This is evident from the prolonged Notch-mediated juxtacrine senescence compared to healthy healing, which means increased production of fibrogenic factors in an untimely manner and delayed transition to an inflammatory SASP. However, this does not lead to a fibrotic state since senescent cell and macrophage derived MMP is able to eventually attenuate ECM levels sufficiently before further progression.

The model simulations presented in this paper clearly show that the differences in the spatiotemporal dynamics of senescent cells lead to several distinct repair outcomes. The difference in senescent cell dynamics can be attributed to differences in SASP composition, duration of senescence and temporal induction of senescence relative to the healing stage. The range of outcomes demonstrated here strongly highlight the dynamic and heterogenous role of senescent cells in wound healing, fibrosis and chronic wounds, and their fine-tuned control. Furthermore, these results suggest that senescent cell manipulation, for example by decreasing the cell population size using senolytics, could be used to reverse unfavourable pathological outcomes, within a treatment window of reversibility.

In summary, the interplay between varying senescent cell activity and their surroundings lead to a wide range of distinct spatiotemporal dynamics, qualitatively summarised in Figure 14: An appropriate number of senescent cells present during the mid to late stages of wound healing promotes healthy healing. However, the presence of excess senescent cells promotes a non-healing chronic wound. Pre-mature induction of senescence or excess pre-existing senescent cells, such as during ageing, also promotes a chronic wound. Insufficient number of senescent cells could result in fibrosis due to lack of ECM breakdown. Lastly, delayed induction of senescent cells could result in fibrosis with delayed fibrolytic activity.

**Figure 14.**
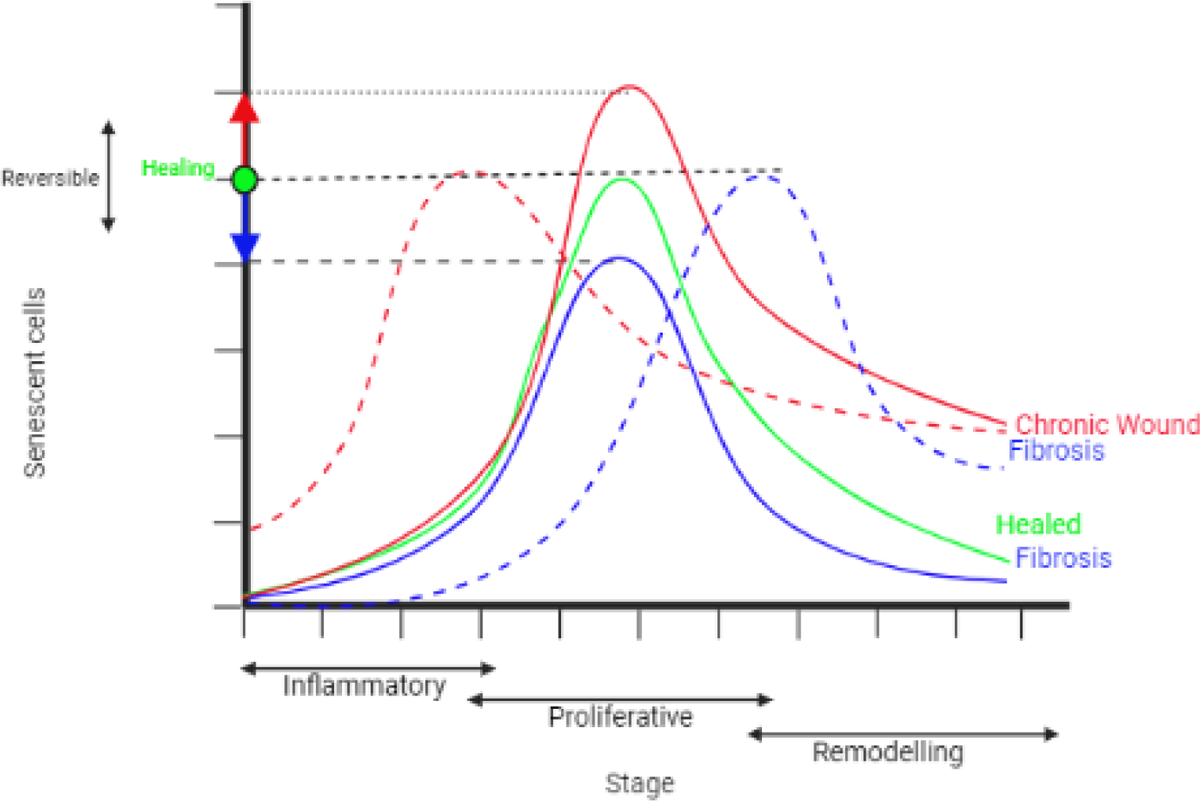
Illustration of the hypothesis for the heterogeneity in senescent cell spatiotemporal dynamics in tissue repair and related pathologies. The wound healing stage is shown on the horizontal axis and senescent cell percentage is shown on the vertical axis. Green solid line: An appropriate number of senescent cells induced during the mid to late stages of wound healing promotes healthy healing. Blue solid line: An insufficient number of senescent cells leads to a fibrotic response. Red solid line: Excessive senescent cells lead to a chronic wound. Red dotted line: pre-mature induction of senescent cells leads to a chronic wound response despite the total percentage of senescent cells being appropriate. Blue dotted line: Delayed induction of senescent cells leads to a fibrotic response, also despite the total percentage of senescent cells being appropriate. Senescent cell dynamics, for example, cell population size, could be manipulated to reverse unfavourable pathological outcomes, within a treatment window of reversibility.

### Limitations and further work

The model is a simplification of highly complex biological processes. Abstraction was necessary to keep the model computationally feasible, while at the same time exploring key processes of interest. The model can potentially be improved by including more immune cell types, different types of ECM components and more growth factors and cytokines that may be of interest. Furthermore, the model was parametrised to existing data to the best of our ability. Using existing data has its pitfalls: firstly, the amount of experimental work (appropriate and relevant for this model) on the dynamics of senescent cells and SASP in healthy and pathological wound healing is limited, and secondly, the data was gathered from different sources and contexts, followed by calibration to the model. While this may not be ideal, this wound healing model can be used as a helpful base to inform experimental work on senescent cell dynamics in healthy and pathological wound healing. This could lead to the development of a more comprehensively parametrised and calibrated computational model, informed by experimental data specific to this context. Several sources of experimental data would be highly valuable to this work, including but not limited to, data on time series cell population changes, cell traction forces, chemical species (such as growth factors, cytokines, chemokines, and proteinases) concentrations, as well as chemical species rate of synthesis and degradation. This would ideally be followed by generating validation data to test the various model predictions. Such a model can be used to study potential treatment avenues, such as senolytics for fibrosis and chronic wounds, and senescent cell manipulation for their beneficial effects. Furthermore, the effect of differences in the duration of the SASP phases (fibrogenic and fibrolytic) and SASP composition on the healing process can also be investigated. This highlights the strength of computational models, as they can easily be updated and improved based on new information and can then be used to inform experimental, and hopefully translational work in a reciprocal manner.

## Acknowledgements

We would like to thank James Wordsworth, Krutik Patel, Rebekah Scanlan and Peter Clark for advice and discussions.

## Author contributions

SC developed the model supervised by DPS with guidance from JPS. SC wrote the paper with input from JPS and DPS.

## Data availability

https://github.com/Sharm8/Senescence_wound_healing_model

